# A mitochondrial redox switch licenses the onset of morphogenesis in animals

**DOI:** 10.1101/2024.10.28.620733

**Authors:** Updip Kahlon, Francesco Dalla Ricca, Saraswathi J. Pillai, Marine Olivetta, Kevin M. Tharp, Li-En Jao, Omaya Dudin, Kent McDonald, Mustafa G. Aydogan

## Abstract

Embryos undergo pre-gastrulation cleavage cycles to generate a critical cell mass before transitioning to morphogenesis. The molecular underpinnings of this transition have traditionally centered on zygotic chromatin remodeling and genome activation^1,2^, as their repression can prevent downstream processes of differentiation and organogenesis. Despite precedents that oxygen depletion can similarly suspend development in early embryos^3–6^, hinting at a pivotal role for oxygen metabolism in this transition, whether there is a *bona fide* chemical switch that licenses the onset of morphogenesis remains unknown. Here we discover that a mitochondrial oxidant acts as a metabolic switch to license the onset of animal morphogenesis. Concomitant with the instatement of mitochondrial membrane potential, we found a burst-like accumulation of mitochondrial superoxide (O_2_^-^) during fly blastoderm formation. *In vivo* chemistry experiments revealed that an electron leak from site III_Qo_ at ETC Complex III is responsible for O_2_^-^ production. Importantly, depleting mitochondrial O_2_^-^ fully mimics anoxic conditions and, like anoxia, induces suspended animation prior to morphogenesis, but not after. Specifically, H_2_O_2_, and not ONOO^-^, NO, or HO•, can single-handedly account for this mtROS-based response. We demonstrate that depleting mitochondrial O_2_^-^ similarly prevents the onset of morphogenetic events in vertebrate embryos and ichthyosporea, close relatives of animals. We postulate that such redox-based metabolic licensing of morphogenesis is an ancient trait of holozoans that couples the availability of oxygen to development, conserved from early-diverging animal relatives to vertebrates.

## Main

Animal embryogenesis starts with proliferative cleavage cycles to build a foundational cell mass in preparation for transitioning to differentiation and organogenesis – a milestone developmental switch that defines the onset of morphogenesis. Seminal genetic approaches paved the way to dissecting the molecular underpinnings of this transition process^7,8^, revealing a set of zygotic molecules produced upon embryonic genome activation. Largely involved in chromatin remodeling and cytoskeletal organization, these factors help establish cell adhesion and polarity at the basis of body patterning and axis-determination^9–11^. As such, a framework centering on zygotic genome activation has emerged for the morphogenetic licensing of early development.

Recent germline-based maternal effect studies, however, paint a more complex picture for the upstream control of this transition, as they reveal an unexpected set of additional gene products related to mitochondrial metabolism, such as protein import^12^, nucleoid maintenance^13^, the tricarboxylic acid (TCA) cycle^14,15^ and oxidative phosphorylation^16^. Although several mitochondria-associated enzymes have been found to gate chromatin remodeling and zygotic genome activation at the onset of morphogenesis^17,18^, exact upstream signals and roles that the mitochondrial metabolism plays in this transition remain largely unknown. Despite reports that such maternal effect mutants can halt development prior to morphogenesis^14,16^, whether there is a *bona fide* metabolic switch that licenses this transition, and if so, how this is achieved chemically remains unclear.

### A mitochondrial ROS burst at the onset of morphogenesis

Classic studies have demonstrated that the absence of oxygen (anoxia) can suspend blastula formation in many animals, from brine shrimps^4,19^ to fishes^5,20^, from worms^6,21^ to fruit flies^3^. Dubbed *suspended animation*, such an early embryonic response to oxygen deprivation has traditionally been attributed to its likely consumption of oxygen for oxidative phosphorylation, as needed to generate ATP in mitochondria. Paradoxically, however, supplying ATP into early embryos incapable of electron transport chain (ETC) or TCA fails to rescue suspended animation^22,23^. Supporting this notion, other works have demonstrated that it is primarily glycolysis that meets the energetic demands of early development in pre-gastrulation invertebrate^24,25^ and vertebrate embryos^26–29^. Together, these results hint at a role for oxygen at the onset of morphogenesis that extends beyond its canonical use in oxidative phosphorylation.

Besides their role in ATP production, mitochondria act as the major source of reactive oxygen species (ROS). As this is another fundamental oxygen-dependent process that serves to signal physiological processes^30,31^, we began by exploring this possibility in *Drosophila* embryogenesis (Figs. 1 and 2). Just as in vertebrates^20^, reaerating anoxic fly embryos can rescue suspended animation without lethality only after, but not before, the onset of morphogenesis^3^. Since this highlights an indispensable role for oxygen during blastoderm formation, we first monitored mitochondrial membrane potential (ΔΨm) in living blastoderms injected with tetramethylrhodamine ethyl ester perchlorate (TMRE) (Fig. 1a). Consistent with the initiation of oxidative phosphorylation at the onset of morphogenesis^24,26–29^, mitochondria gradually instated their ΔΨm starting largely from cycle 12 during blastoderm formation and plateauing in cycle 14 (Fig. 1b). Surprisingly, however, in embryos treated with CellROX – a probe approximating superoxide (O_2_^-^) formation^32^ – we also observed a burst of ROS production that accompanies ΔΨm instatement at the onset of morphogenesis (Fig. 2a,b).

**Fig. 1:**
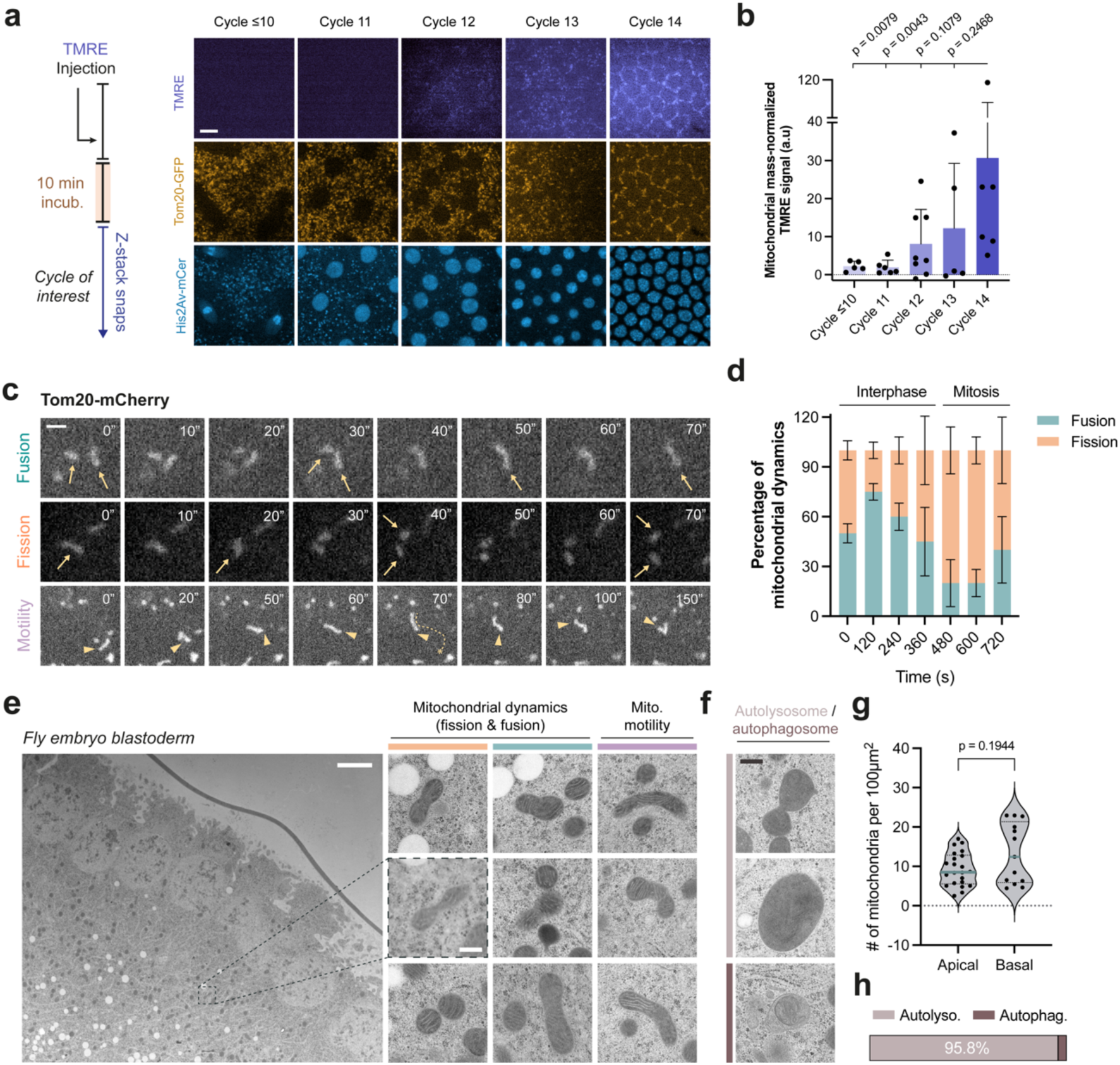
Embryonic mitochondria activate during blastoderm formation in flies. **a**, Confocal micrographs depicting the status of blastoderm mitochondrial ΔΨm in embryos expressing His2Av-mCer and Gal4^V37^>UAS-Tom20-GFP injected with the TMRE dye. Sketch (left) illustrates the experimental protocol (see *Methods*). **b**, Bar graph quantifying the mitochondrial mass-normalized TMRE signal from (a) in cycle 10 (n=5), 11 (n=6), 12 (n=8), 13 (n=5) and 14 (n=6) embryos. Data (mean ± s.d.) are compared with two-tailed Mann-Whitney tests. **c**, Confocal micrographs demonstrating representative mitochondrial fusion, fission and motility events in unperturbed embryos expressing Gal4^V37^>UAS-Tom20-mCherry. Arrows direct attention to the persistence of fused and fragmented mitochondria over time, as demonstrated. Arrowhead is fixed to the same tip of a mitochondrion as it moves, demonstrating its motility. **d**, Stacked bar graph quantifying the percentage of fusion or fission incidents observed for mitochondria that display membrane dynamics during cycle 12 (n= 20 mitochondria per time point, except n=10 at t=720s). Data (mean ± s.e.m.) were plotted to represent an average of mitochondrial behaviour from multiple embryos (see Extended Fig. 1c for individual embryos). **e**, Electron micrographs (EM) depicting the blastoderm of an unperturbed fly embryo with insets demonstrating signatures of membrane dynamics and motility on individual mitochondria, as assessed by previously established EM ultrastructure guidelines (see *Methods*) and in line with observations in (c). **f**, EM insets demonstrating the existence of autophagosomes and autolysosomes in fly embryos. **g**, Violin plots comparing the number of mitochondria across the dorso-ventral axis of the fly blastoderm as observed in 100µm^2^ sized apical and basal EM micrographs (n=22 and 13 sections respectively). Data points represent individual sections, whose distributions are indicated with quartile lines and a probability density estimation using the kernel plot. Apico-basal difference was compared with a two-tailed Mann-Whitney test. **h**, Lateral stacked bar comparing the percentage of autolysosomes vs. autophagosomes observed among all macroautophagy events (n=48 encountered scanning through 13,560µm^2^ worth EM sections). Scale bars, 5µm (**a**,**e**), 1µm (**c**), 0.5µm (insets in **e**,**f**)

**Fig. 2:**
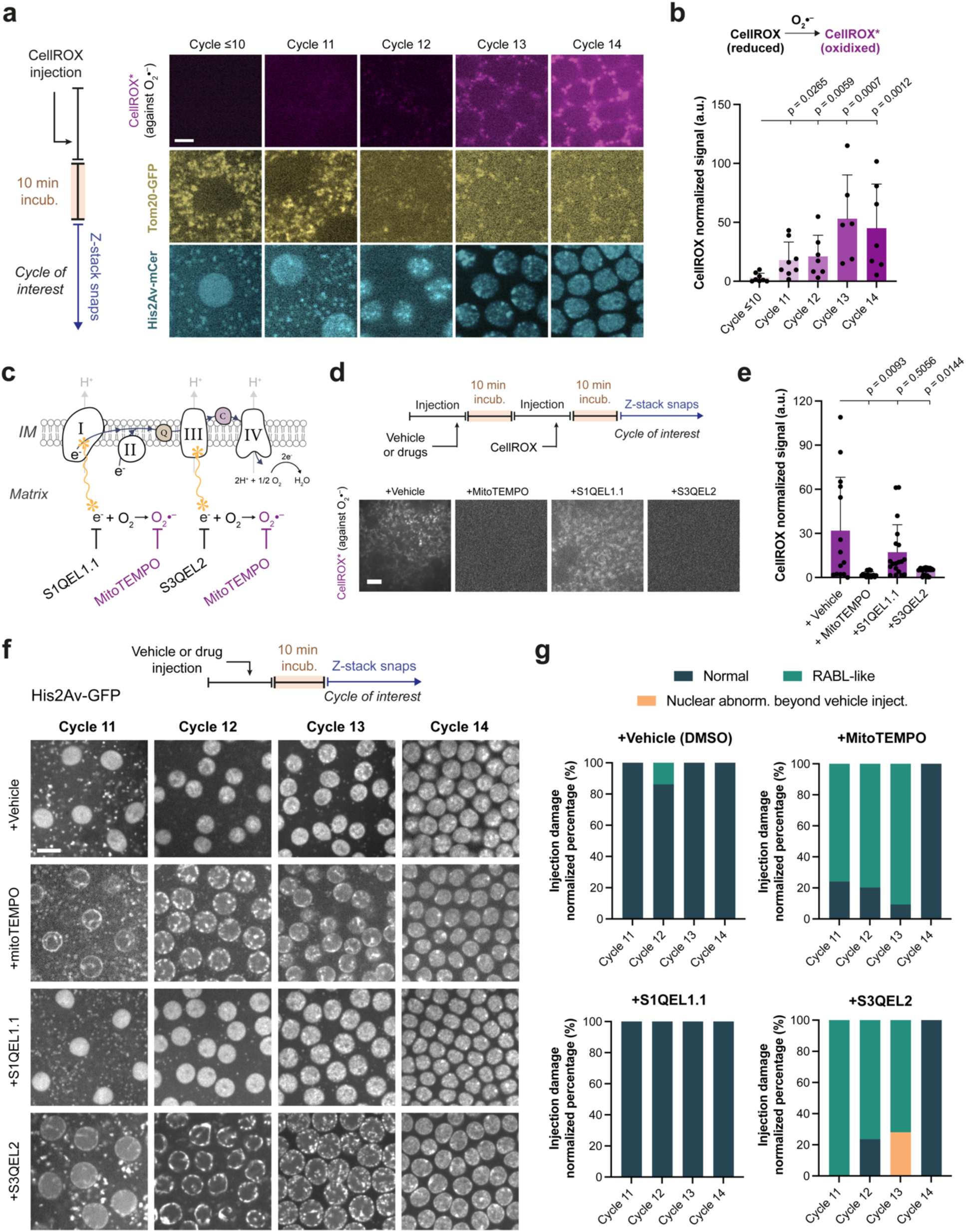
A mitochondrial ROS (mtROS) licenses the onset of morphogenesis. **a,** Micrographs depicting a burst-like increase of O_2_^-^ during blastoderm formation in embryos expressing His2Av- mCer and Gal4^V37^>UAS-Tom20-GFP injected with the CellROX probe. **b**, Bar graph quantifying the mitochondrial mass-normalized CellROX signal from (a) in cycle 10 (n=8), 11 (n=9), 12 (n=7), 13 (n=6) and 14 (n=7) embryos. Data (mean ± s.d.) are compared with two-tailed Mann-Whitney tests. **c**, Cartoon depicting pharmaceutical strategies to prevent e^-^ leak at prominent ETC sites that lead to O_2_^-^ production. **d**, Micrographs depicting CellROX probe signal in embryos administrated with indicated treatments in prior (see Extended Data Fig. 3 for all conditions across different blastoderm cycles). **e**, Bar graph quantifying the mitochondrial mass-normalized CellROX signal from the experiments (d) in embryos pre-treated with vehicle (n=14), MitoTEMPO (n=17), S1QEL1.1 (n=19), and S3QEL2 (n=27) conditions. Data (mean ± s.d.) are compared using a Welch’s t test (for Gaussian distributed data) or a Mann-Whitney test. **f**, Micrographs demonstrating representative nuclear/chromatin phenotypes under the indicated conditions at cycles 11-14 in embryos expressing His2Av-GFP. **g**, Bar graphs quantifying the nuclear/chromatin configuration phenotypes observed under the conditions signified in (f). To ease comparison between the normal vs Rabl-like conditions, the percentages presented here were normalized to exclude several otherwise karyotype damages induced via the vehicle injection itself by default. As such, see Extended Data Fig. 6d-g for associated sample (embryo) numbers. Sketches in (**a**, **d**, and **f**) illustrate the experimental protocols (see *Methods*). Scale bars, 3µm (**a**), 5µm (**d**), 8µm (**f**).

Excess ROS production can damage mitochondrial motility^33^ and dynamics^34^, leading to their elimination by fragmentation and mitophagy^35^. Given that embryos have an inherent capacity to induce mitophagy – as observed in allophagy during paternal mitochondrial elimination^36–38^ – we tested whether the ROS burst during blastoderm formation could lead to similar mitochondrial responses. In embryos expressing Tom20-mCherry (a mitochondrial outer membrane marker; Supplementary Video 1), we found that the blastoderm mitochondria are dynamic and go through fission/fusion (Fig. 1c) despite the ROS burst and their relatively small sizes (Extended Data Fig. 1a). Just as their counterparts in somatic tissues^39^, mitochondrial fission-fusion cycles appeared to be in sync with the progression of the cell cycle (Fig. 1d and Extended Data Fig. 1b,c), in which mitochondria mostly fuse during interphase and go under fission during mitosis. We found that fusing mitochondria was on average smaller than those that undergo fission (Extended Data Fig. 1a,d), implicating that the regulatory mechanisms responsible for mitochondrial size and morphology homeostasis^40^ are intact in these embryos. Based on previously established ultrastructure guidelines (see *Methods*), we also confirmed these by electron microscopy, examining signatures of mitochondrial morphology in sectioned wild-type embryos (Fig. 1e-h). Since mitochondria were homogenously spread across the apico-basal axis (Fig. 1g), we concluded that mitochondrial motility continues to serve its function^41^ of homogenously distributing these organelles at this stage of development. As evident from most autophagosomes filled with undigested lipids in the form of autolysosomes (Fig. 1f; see *Methods*), we observed active autophagy during fly blastoderm formation, yet never found any instances of mitophagy (Fig. 1h).

Given that blastoderm mitochondria are dynamic and appear to be regulated normally, these results suggested that the burst of ROS production (Fig. 2a,b) is unlikely to be an inadvertent consequence of mitochondrial activation at this stage of development. As such, we next aimed to identify the exact sources of embryonic ROS, enabling us to investigate its potential role at the onset of morphogenesis.

### Mitochondrial Complex III mediates the ROS production at the onset of morphogenesis

Injecting CellROX into embryos expressing Tom20-GFP, we found that most ROS production appears to co-localize with mitochondria (Extended Data Fig. 2a,b). To directly test whether mitochondria can account for embryonic ROS production, we examined CellROX in embryos pre-treated with MitoTEMPO, a mitochondria-targeted antioxidant that specifically scavenges O_2_^-^ (Fig. 2c). We found that mitochondria can indeed fully account for the CellROX signal we observe in these embryos (Fig. 2d,e and Extended Data Fig. 3a,b).

Mitochondrial O_2_^-^ is generated by electrons (e^-^) prematurely leaking from e^-^ carriers through ETC and reacting with oxygen. Although there are at least eleven reported sites of leak in mitochondria^42^, growing evidence highlights the ubiquinone-reducing site of Complex I (site I_Q_) and the ubiquinol-oxidizing site of Complex III (site III_Qo_) as the primary leak sources for O_2_^-^ as relevant to physiology^43,44^ (Fig. 2c). To test whether e^-^ leak from ETC can account for the mitochondrial O_2_^-^ production, we leveraged a new class of small molecule suppressors that selectively eliminate O_2_^-^ production by Complex I at site I_Q_ (S1QEL1.1) and Complex III at site III_Qo_ (S3QEL2) without affecting the e^-^ transfer or altering oxidative phosphorylation^45,46^ (Fig. 2c). Although targeting e^-^ leak at Complex I appeared to mildly influence O_2_^-^ production in mitochondria, the effect was not statistically significant (Fig. 2d,e and Extended Data Fig. 3c). Targeting the e^-^ leak at Complex III, however, completely abolished mitochondrial O_2_^-^ production and fully mimicked the results with MitoTEMPO treatments (Fig. 2d,e and Extended Data Fig. 3b,d). Together, these results indicate that e^-^ leak at Complex III site III_Qo_ is the primary source of mitochondrial O_2_^-^ at the onset of fly morphogenesis.

### Complex III-mediated ROS production acts as a metabolic switch to license the onset of morphogenesis

Even a normally functioning ETC can leak electrons and produce O_2_^-^ at basal levels, so the mitochondrial O_2_^-^ that we observe (Fig. 2a and Extended Data Fig. 2 and 3) could simply be a byproduct of ETC activation. Since ROS formation is restricted by oxygen availability, we sought to test whether preventing O_2_^-^ production would functionally mimic the suspended animation induced by anoxia during blastoderm formation^3^ (Fig. 2f). Remarkably, both scavenging mitochondrial O_2_^-^ altogether and eliminating e^-^ leak specifically at Complex III (site III_Qo_) suspended embryogenesis by a developmental arrest with chromatin condensed in a Rabl-like configuration – the exact cytological response that anoxia uniquely elicits during the embryogenesis of flies^3^ and other invertebrates^47^ (Fig. 2f and Extended Data Fig. 4; see how this arrest is robust even to exogenous perturbations in Supplementary Video 2). Reminiscent of the switch-like viability difference of anoxic embryos with reference to the onset of morphogenesis^3,20^, both types of O_2_^-^ elimination induced suspended animation only during the blastoderm formation (nuclear cycles 11-13) preceding the onset of morphogenesis, but not after (cycle 14) (Fig. 2f,g and Extended Data Fig. 4).

Suspended animation of anoxic embryos necessitates a halt to the nuclear division cycle^3,5,6^ – just as we also find is the case for oxidant-depleted embryos (Fig. 2f and Supplementary Video 2). Although redox-based activation of cell cycle regulators^48,49^ can explain how such a nuclear arrest can be achieved, emerging evidence indicates that aspects of cytoplasmic organization, divisions and even differentiation can occur autonomously of the mitotic CDK/cyclin complexes and nuclear divisions in early development^50–55^. Since lack of oxidant signaling can similarly impair myosin recruitment to plasma membrane^56–58^ and actin polymerization^59,60^ – two key drivers of cortical contractility during gastrulation – we tested whether eliminating O_2_^-^ production prior to morphogenesis can also impair components of cortical contractility beyond a nuclear cycle arrest. To this end, we examined the behavior of non-muscle myosin II (by MRLC-GFP) and actin (by Moe-ABD-GFP) upon suspended animation. We found that myosin recruitment to the plasma membrane and cortical actin organization were abolished in a strikingly all-or-none manner (Extended Data Fig. 5), directly mimicking the inhibition of Rho-kinase activity and actin polymerization in these embryos^53^. As such, distinguishing it from a canonical cell cycle arrest that cannot prevent cortical organization needed for gastrulation^54,61^, eliminating O_2_^-^ during blastoderm formation prevents progression into morphogenesis in part by halting cortical organization.

Together, these results demonstrate that mitochondrial Complex III-mediated ROS production in early embryos normally acts as a ‘metabolic switch’ to license the onset of fly morphogenesis.

### HO•, NO or ONOO^-^ cannot account for the mtROS-based switch

O_2_^-^ molecules are the most upstream ROS produced in mitochondria. However, these oxidants are short-lived^62^ (as minimally as t_1/2_ of milliseconds), as they are almost simultaneously converted to more stable oxygen and nitrogen species upon their production (Fig. 3a). Since a capacity to act as a signaling molecule would require stability to diffuse longer distances, we conjectured that a downstream oxidant species must act as a key signal for the mtROS-based switch in embryos. We first investigated hydroxyl radicals (HO•) (Fig. 3a). To test whether their depletion can induce the Rabl-like phenotype, we leveraged dimethyl sulfoxide (DMSO), an organosulfur compound that – at 5×10^-2^ to 1M *in vitro* – fully scavenges HO• ^63,64^. Factoring in ∼20x dilution upon microinjections, we examined embryos expressing fluorescent His2Av (chromatin) using a dilution series of DMSO with a maximum needle concentration at 14M (Extended Data Fig. 6a-d). As an injection control, we used water to account for their injection-induced repercussions (e.g., telophase defects and chromosome damages). Fortuitously, the DMSO experiments also acted as a vehicle control per its use as a solvent in our experiments to eliminate mitochondrial O_2_^-^ production (Extended Data Fig. 6e-g). Across the dilution series, and even at the highest possible [DMSO], we found that majority of the embryos and their nuclei appeared relatively normal (Extended Data Fig. 6a-d), and rest of the phenotypes largely overlapped with the defects observed by the injection procedure itself with water (Extended Data Fig. 6b).

**Fig. 3:**
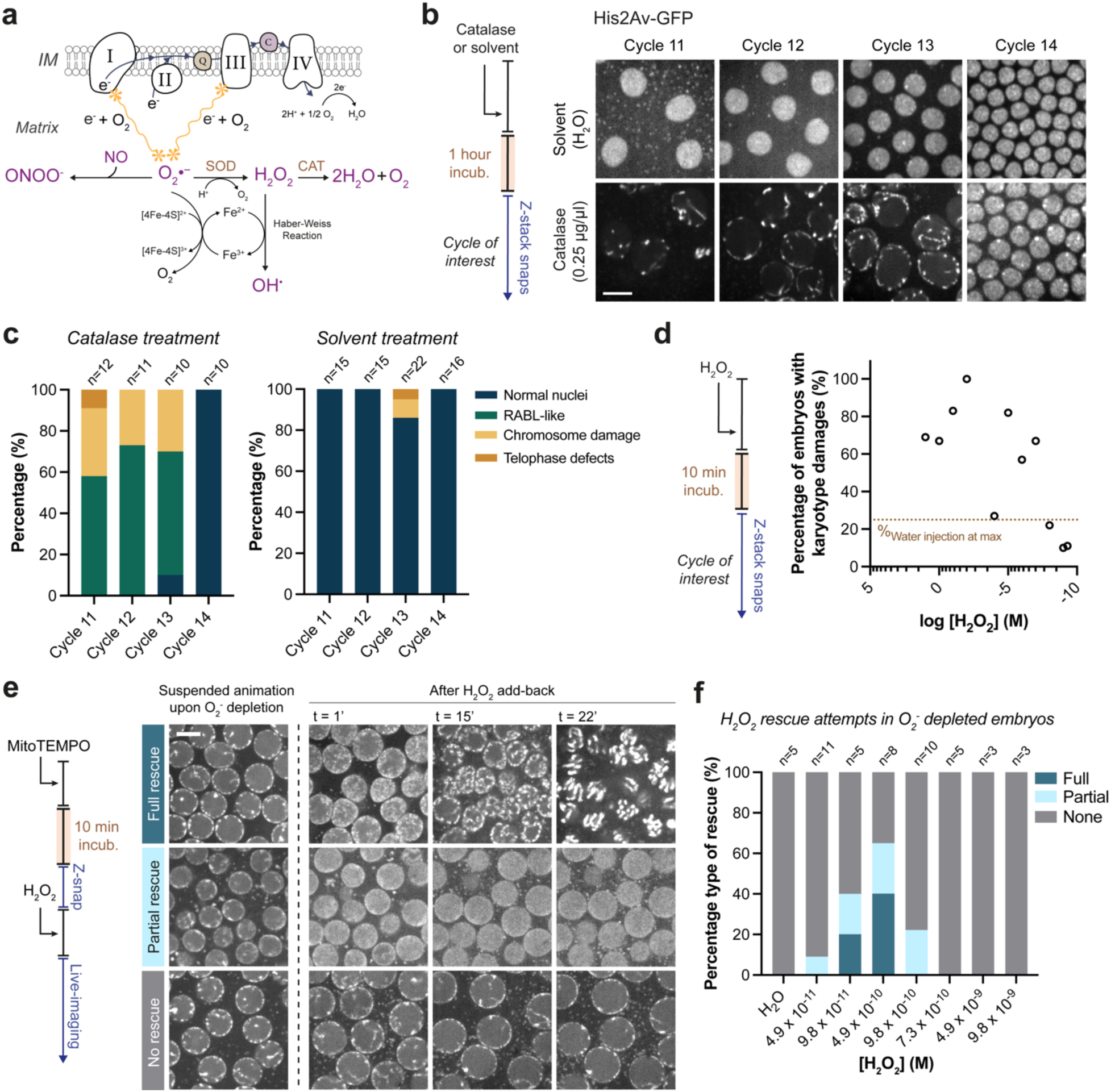
H_2_O_2_ molecules are necessary and sufficient for the mtROS-based switch at the onset of morphogenesis. **a**, Cartoon depicting the chemical landscape of reactive oxygen and nitrogen species stemming from O_2_^-^ produced via e^-^ leak through ETC. **b**, Micrographs demonstrating representative nuclear/chromatin phenotypes at cycles 11-14 in His2Av-GFP embryos injected with solvent vs. catalase. **c**, Bar graphs quantifying the nuclear/chromatin configuration phenotypes observed under the conditions signified in (b) with the associated sample (embryo) numbers. **d**, Scatter plot quantifying the percentage of embryos with karyotype damages (as depicted in Extended Data Fig. 10) as a function of [H_2_O_2_] with which His2Av-GFP embryos were treated. *Brown* dotted line represent the maximum level of specimen percentile with karyotype damages even when injected with water. **e**, Micrographs depicting the three distinct phenotypes observed in His2Av-GFP embryos that were initially treated with MitoTEMPO to induce suspended animation and subsequently were injected with various concentrations of H_2_O_2_ to rescue from this developmental arrest. *Full rescue* signifies embryos that resumed to their mitotic cycling, whereas *Partial rescue* indicates embryos whose nuclei decondensed from the Rabl-like state yet did not resume mitotic cycling. **f**, Stacked bar graph quantifying the phenotypes from (e) in MitoTEMPO-arrested embryos that were attempted to rescue with a dilution series of physiological [H_2_O_2_], as determined by the experiments in (d). Associated sample (embryo) numbers are indicated above each stacked bar. Sketches in (**b**, **d**, and **e**) illustrate the experimental protocols (see *Methods*). Scale bars, 10µm (**b**), 5µm (**e**).

As a potent hygroscopic agent inducing dehydration near lipid membrane surfaces^65^, DMSO also induced shrinking of nuclei, making them appear as though they are desiccated. Interestingly, early embryos appeared particularly vulnerable to such dehydration (Extended Data Fig. 6d). We currently do not know the exact reasons for this vulnerability, but because early embryos are concentrated with unusual amounts of maternally deposited lipids and carbohydrates^25,66,67^, they may be particularly sensitive to dehydration. Indeed, we never observed this phenomenon during water injections (Extended Data Fig. 6a,b), even when performed systematically across different cycles (Extended Data Fig. 8f). Meanwhile, we almost never encountered a Rabl-like phenotype during our DMSO experiments (n=92 embryos) – except for one embryo, which accounts for only ∼2% of the total number of injections even at the highest possible [DMSO]. We ascribe this outlier to the vast feedback networks that regulate redox chemistry *in vivo*: depleting one type of free radical can influence the production of another, as exemplified by the case of O_2_^-^ molecules *vide infra* (Extended Data Figs. 7 and 8). Together, these suggested that HO• is unlikely to account for the mtROS-based switch at the onset of morphogenesis.

We next tested the involvement of nitric oxide (NO) and peroxynitrite (ONOO^-^), two nitrogen species that are intricately linked with mitochondrial ROS metabolism^69^ (Fig. 3a and Extended Data Fig. 7a). Although, NO signaling has been previously studied in fly embryos using exogenous supplement of its donors^22^ (e.g., S-Nitroso-N-acetylpenicillamine), whether it is generated endogenously in this system remains unknown. Since ONOO^-^ is produced by the reaction of NO and O_2_^-^, we first probed ONOO^-^ with a selective fluorescent probe conjugated to a hydrazide group^70^, acting also as a simultaneous indicator for the presence of endogenous NO (Extended Data Fig. 7a). Consistent with its affinity to induce lipid peroxidation and to oxidize low-density lipoproteins^71,72^, we found that endogenous ONOO^-^ was largely associated with yolk granules in the embryo interior (basal) and appeared diffusely in the blastoderm cytosol (apical) (Extended Data Fig. 7b). Interestingly, the chromatin in embryos injected with the ONOO^-^ fluorescent probe occasionally displayed the Rabl-like phenotype (Extended Data Fig. 8a,b). As this probe fluoresces by *reacting* with ONOO^-^ molecules (Extended Data Fig. 7a), we suspected that diminishing ONOO^-^ levels by this reaction may trigger suspended animation. However, it is also possible that such prolonged ONOO^-^ depletion could impair feedback mechanisms that normally prevent excess NO production via the availability of ONOO^-^ itself^72^, resulting in an amplified sink of O_2_^-^ due to its increased consumption by NO. To resolve between these possibilities, we reasoned to prevent ONOO^-^ formation by scavenging NO, as the latter’s production is a limiting factor for the former independently of O_2_^-^ (see Extended Data Fig. 7a). Leveraging a widely used NO scavenger to perturb ONOO^-^ production (Extended Data Figs. 7c,d), we found that this is indeed the case: neither decreasing ONOO^-^ levels nor diminishing NO is sufficient to trigger the Rabl-like phenotype in a significant way (Extended Data Fig. 8c-f).

In summary, our experiments suggested that – like the HO• – depleting NO and ONOO^-^ alone cannot account for the Rabl-like phenotype. These results are also consistent with the dispensability of nitric oxide synthase for *Drosophila* development^73^. Instead, another oxidant that stem from O_2_^-^ must be responsible for the mtROS-based switch at the onset of morphogenesis.

### H_2_O_2_ molecules are single-handedly necessary and sufficient for the mtROS-based switch

We finally investigated hydrogen peroxide (H_2_O_2_). First, we assessed whether the mitochondrial O_2_^-^ production is accompanied by H_2_O_2_ formation during blastoderm formation, as would be expected from their simultaneous dismutation by superoxide dismutase (Extended Data Fig. 9a). Using MitoPY1, a boronate-based probe that fluoresces upon its selective reaction with H_2_O_2_ in mitochondria^75^, we found that this is indeed the case (Extended Data Fig. 9b,c). Surprisingly, the injected embryos occasionally displayed the Rabl-like phenotype (Extended Data Fig. 9d,e). As MitoPY1 fluoresces by *reacting* with H_2_O_2_ molecules and may lead to its partial depletion (Extended Data Fig. 9a), we sought to test whether a more complete depletion of H_2_O_2_ by its catalysis can consistently suspend embryogenesis. As catalase breaks down H_2_O_2_ into water and oxygen (Fig. 3a), we administrated this oxidoreductase (purified from bovine liver) and examined its effects in embryos expressing His2Av-GFP. Remarkably, the catalytic decomposition of H_2_O_2_ halted embryonic development in a Rabl-like arrest (Fig. 3b,c). As such, we conclude that H_2_O_2_ production is necessary to satisfy the mtROS-based switch.

To test whether H_2_O_2_ can be sufficient for the mtROS-based switch, we sought to perform “rescue” experiments. Upon a cumulative depletion of mitochondrial ROS, we asked whether adding back only H_2_O_2_ could release embryos from suspended animation. By taking advantage of H_2_O_2_’s widely observed karyotypic toxicity at supraphysiological concentrations^76–79^, we first determined a range of add-back [H_2_O_2_] needed *in vivo*. Across a dilution series starting with off-the-shelf concentration at 9.8M, we examined the effects of H_2_O_2_ in embryos that express His2Av-GFP (Extended Data Fig. 10). As expected, a palette of karyotype damages ensued across the dilution series (Extended Data Fig. 10a; see *Methods* for the phenotype nuances), which gradually declined as [H_2_O_2_] was lowered (Extended Data Fig. 10b). With water injection-induced damages as the benchmark, we established a “physiological” range for the add-back [H_2_O_2_] needed for our prospective rescue experiments (Fig. 3d). Next, we pre-treated blastoderm-stage embryos with MitoTEMPO to induce suspended animation (Fig. 3e). We then attempted to rescue the suspended embryos by adding back H_2_O_2_ (Fig. 3e,f). Remarkably, we achieved a complete rescue in 40% of our trials at a concentration of 4.9×10^-10^ M (Fig. 3e,g and Supplementary Video 3; dubbed ‘full’ rescue), in which chromatin decondensed from the Rabl-like configuration and resumed its mitotic progression, albeit with telophase errors (Supplementary Video 3). As a control, we added back water and did not observe a rescue (Fig. 3f and Extended Data Fig. 11d,e).

Within the concentration range of 4.9×10^-11^ to 9.8×10^-10^ M, resupplying H_2_O_2_ also led to an incomplete rescue phenotype (dubbed ‘partial rescue’), in which chromatin decondensed yet did not resume the cell cycle (Fig. 3e,g and Supplementary Video 4). Surprisingly, at concentration ranges closer to supraphysiological conditions, adding back H_2_O_2_ did not induce either of the rescue phenotypes (Fig. 3e,f and Supplementary Video 2). We hypothesize that chromatin damages induced by excess H_2_O_2_ may also impair nuclear ability to go through the mitotic cycle, hence overriding H_2_O_2_’s ability to fully rescue suspended animation.

Could suspended animation in oxidant-depleted embryos somehow stem from a loss of overall capacity to generate ATP? We tested this possibility by mimicking the above experiments for H_2_O_2_, but by adding back ATP. It is well established that excess ATP chelates Mg^+2^^79–81^ – a divalent cation critical for the dynamic instability of microtubules^83–85^, hence required for proper spindle assembly and chromosome segregation. As such, we aimed to determine an optimal add-back concentration by examining the karyotypic toxicity of ATP at 0.5–5mM, the homeostatic range of intracellular ATP^86^ (Extended Data Fig. 11). Accounting for ∼20x dilution from a maximum needle concentration at 100mM (i.e., ∼5mM effective conc.), ATP injections – even across this homeostatic range – were largely associated with a series of mitotic defects (Extended Data Fig. 11a). Only near the range minima (∼0.6mM effective conc.), the injected embryos developed normally (Extended Data Fig. 11b,c). Using this concentration, we attempted to rescue MitoTEMPO-treated embryos from suspended animation with ATP, but to no avail (Extended Data Fig. 11d,e). Like previous studies that failed to rescue early embryos with impaired of ETC or TCA using ATP^22,23^, we conclude that the same holds true for suspended animation triggered by oxidant depletion, even at the highest tolerable [ATP] supplement.

Collectively, these results demonstrate that H_2_O_2_ is necessary and sufficient for the mtROS-based switch at the onset of morphogenesis. Since simply decondensing chromatin is not sufficient to enable mitotic cycling (Fig. 3e and Supplementary Video 4), H_2_O_2_’s role to satisfy the mtROS- based switch in oxidant-depleted embryos appears *at least* two-fold: first by releasing chromatin from its Rabl-like configuration, then by permitting cell cycle progression. These findings highlight the complexity of the mtROS-based switch, paving the way towards identifying the genetic and biochemical basis of metabolic licensing in early animal development.

### mtROS-based oxidant switch is conserved from animal relatives to vertebrates

Oxygen depletion-induced suspended animation is a widely conserved phenomenon in the early embryogenesis of many animals^3–6,19^ (Fig. 4a). Such capacity to induce hypometabolism may possibly be a common feature in the early life cycles of opisthokonts more generally, as oxygen depletion can suspend even the sporulation of *Saccharomyces cerevisiae*^87^. Thus, we sought to explore whether the mtROS-based switch that we observe at the onset of early invertebrate development could be a hallmark of licensing morphogenesis metabolically across holozoans.

**Fig. 4:**
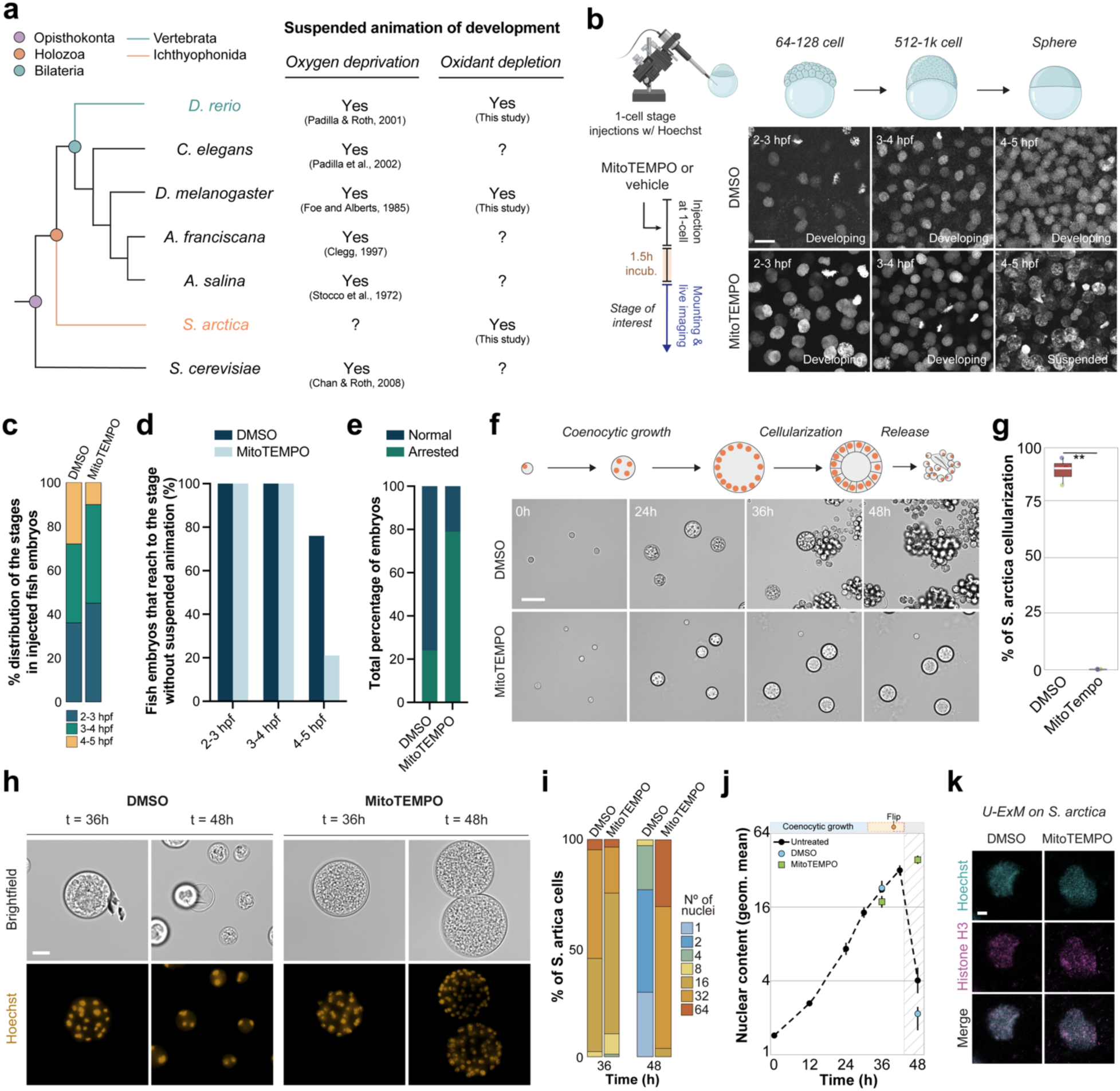
mtROS-based licensing of morphogenesis is conserved from relatives of animals to vertebrates. **a**, Cladogram highlighting the position of *Drosophila melanogaster* (*D. melanogaster*) among other commonly explored animals and species in the broader clade of opisthokonts (e.g. fungi) in anoxia studies. Whether early morphogenetic events in the life cycles of these species respond to oxygen deprivation and/or oxidant depletion by a suspended animation (or an alike state) are indicated next to taxonomic names with their key citations. *Danio rerio* (*D. rerio*) in *light teal* and *Sphaeroforma arctica* (*S. arctica*) in *orange* are highlighted as key organisms that could offer insights into the conservation and evolution of the mtROS-based switch in animals and their close holozoan relatives. **b**, Maximum intensity-projected micrograph stacks depicting the progression of 2-4 hours post-fertilization (hpf) cleavage division cycles in *D. rerio* embryos injected with either DMSO (vehicle) or MitoTEMPO at 1-cell stage, while bathing in medium containing Hoechst dye (marking chromatin). Note the suspended animation at ∼4hpf under MitoTEMPO condition. Sketch (left) illustrates the experimental protocol (see *Methods*). **c**, Stacked bars quantifying the percentage distribution of cleavage division stages observed during the experiments in (b), from 4 independent trials for both DMSO (n=25 embryos) and MitoTEMPO (n=24 embryos). Note the decrease of 4-5hpf zebrafish embryos under MitoTEMPO conditions, reflecting the suspended animation that takes place between 2-3hpf and 4-5hpf of early development. **d**, Bar graphs quantifying the fraction of zebrafish embryos that can survive to 4- 5hpf of early development without suspended animation under DMSO vs. MitoTEMPO conditions. **e**, Stacked bars quantifying the percentage of arrested embryos under DMSO vs. MitoTEMPO conditions at 4-5hpf of early zebrafish development. **f**, Time-lapse micrographs displaying the life-cycle of *S. arctica* starting from t=0h bathed in either DMSO or MitoTEMPO. Note how MitoTEMPO treatment prevents cellularization and release of *S. arctica*. **g**, Box plots comparing the percentage of *S. arctica* undergoing cellularization bathed in either DMSO or MitoTEMPO (n>213 cells per condition from 3 independent trials). Data (mean ± s.d.) are compared with the t-test (**p<0.005). **h**, Micrographs demonstrating nuclear content (as stained by Hoechst) in *S. arctica* at 36h vs. 48h after 1-nucleus stage under DMSO vs. MitoTEMPO conditions. Note how *S. arctica* can complete its cellularization and release at 48h bathed in DMSO but not in MitoTEMPO. **i**, Stacked bars quantifying the distribution of nuclear content from the experiments in (h), from 3 independent trials for each condition (n>30 cells per time point). **j**, Graph quantifying the mean nuclear content (expressed as log2 of geometric mean) in *S. arctica* at 36h vs. 48h after 1-nucleus stage under DMSO vs. MitoTEMPO conditions. n>213 cells per condition from 3 independent trials. *Black* dotted line represents the nuclear content trajectory of untreated *S. arctica* (data from Olivetta et al.^93^). Data are represented as mean ± s.d. **k**, Chromatin architecture of *S. arctica* treated with either DMSO or MitoTEMPO, then fixed and stained with Hoechst and anti-histone H3 antibody via an ultrastructure expansion microscopy protocol (U-ExM) (see *Methods*). Cartoon diagrams of (**b**) zebrafish embryos and (**f**) *S. arctica* depict developmental or life-cycle stages pertinent to the experiments. Scale bars, 20µm (**b**), 50µm (**f**), 25µm (**h**), 5µm (**k**).

To test the involvement of an mtROS-based switch at the onset of vertebrate morphogenesis, we used early *Danio rerio* (zebrafish) embryos – a model vertebrate for oxygen depletion studies^5,20,88^. Using MitoTEMPO to scavenge mitochondrial O_2_^-^, we examined the progression towards blastula formation. Remarkably, embryos injected at the 1-cell stage suspended their cell cycles only after 1k cell stage (around oblong-sphere stages) during blastula formation prior to the onset of morphogenesis (Fig. 4b-e, Extended Data Fig. 12, and Supplementary Video 5). Just as in fly embryos (Extended Data Fig. 4), we examined whether preventing O_2_^-^ production in fish embryos would phenotypically mimic the suspended animation induced by oxygen depletion in this system^5^. As embodied by a chromatin condensation phenotype in a pre-mitotic/prophase arrest^5^, we found that this is indeed the case (Extended Data Fig. 12).

Inspired by the strict requirement for oxygen in fungal sporulation^87^ (Fig. 4a), we next explored how widely such an mtROS-based switch could be conserved across holozans. To this end, we examined ichthyosporeans, protists positioned between animals and fungi, and considered among the closest relatives of animals with choanoflagellates and filastereans. Leading to the emergence of animals, ichthyosporeans are speculated as key species towards the evolution of embryogenesis^89,90^. Under active scrutiny, this is not least because their morphogenesis succeeds the formation of a multicellular palintomic^90,91^, or multinucleate coenocytic^92,93^, epithelial structure that exhibits similarities and behaves like the animal blastulae and blastoderms respectively^93,94^. In *Sphaeroforma arctica*, a coenocytic ichthyosporean, the assembly of a blastoderm-like structure is followed by a cellularization-based morphogenesis event^92^ (Extended Data Fig. 13) – reminiscent of fly embryos^89^. As such, we tested whether eliminating mitochondrial O_2_^-^ influences the progression of an initially single-celled *S. arctica* to its transient multicellular layer before its redispersal (Fig. 4f). Although *S. arctica* cultured in medium containing MitoTEMPO grew very similarly to their control counterpart, they strikingly suspended development at the onset of morphogenesis immediately prior to cellularization (Fig. 4g-j and Supplementary Video 6). Unlike fishes and flies, however, the chromatin of *S. arctica* was not associated with a particular chromosome condensation phenotype during suspended animation (Fig. 4h), even when we assessed chromatin configuration by ultrastructure expansion microscopy (Fig. 4k). Given that *S. arctica* lack proteins such as BAF and KASH critical for tethering chromatin to the nuclear envelope^90^, these results hint at the emergence of novel cytological features associated with suspended animation towards the evolution of metazoans.

Together, the exceptional similarities we observe among oxidant-depleted ichthyosporeans and the embryos of flies and fishes compel us to postulate that an mtROS-based metabolic licensing of morphogenesis is an ancient trait of holozoans, from close animal relatives to vertebrates.

## Discussion

Despite early proposals by pioneers Jacques Loeb and Joseph Needham on the possibility of a chemical basis for licensing morphogenesis^95,96^, whether there is a *bona fide* switch that controls this in development, and if so, how it could be achieved molecularly remain unknown. Here we discover in flies that mitochondrial Complex III-mediated ROS production acts as a metabolic switch to license the onset of morphogenesis. Our findings from fishes and close relatives of animals suggest the striking possibility of a broad conservation of this switch across holozoans.

By elucidating the function of oxygen in early embryogenesis, evidence of an mtROS-based switch licensing morphogenesis resolves several paradoxes of classic studies that established the indispensability of oxygen in animal development. For instance, anoxia-induced suspended animation during blastoderm formation has been ascribed to a hypometabolic state that emerges from perturbing mitochondrial ATP production^3^. Yet, it has remained enigmatic why halting oxidative phosphorylation (e.g. by hypoxia^22^ or mutations against the ATP synthase machinery^16^) fails to trigger the Rabl-like arrest observed under anoxic conditions. In fact, these perturbations trigger a metaphase arrest and/or defects associated with chromosome segregation^16,22^, consistent with a partial manipulation of ATP levels as the same studies report^16,22^ and as found in our experiments (Extended Data Fig. 11). Combined with reports of glycolysis as the primary energy source in pre-gastrulation invertebrate^24,25^ and vertebrate embryos^26–29^, our results attribute a new role for oxygen prior to the onset of morphogenesis. This can also explain, at least in part, why providing embryos incapable of ETC^22^ or TCA^23^ with ATP is not sufficient to rescue suspended development. As ΔΨm fully reinstates only by cycle 13-to-14 transition in fly embryos (Fig. 1a,b), our results may also explain why maternal effect mutations against the ATP synthase machinery can ramify a phenotype only by these cycles during fly development^16^. Furthermore, as both mitochondrial ATP and ROS production source electrons from ETC, our results may provide a new framework for interpreting why ETC inhibition might prevent developmental progression at similar stages of animal embryogenesis more generally^97^.

To this end, recent studies have demonstrated that oocytes, which are under ‘respiratory quiescence’, become more oxidative upon fertilization in various animals^98–100^. An activity-based profiling of H_2_O_2_-modified cysteines has revealed that reactive thiols are largely enriched in proteins associated with mitotic processes during fly oocyte-to-embryo transition^99^. However, it remains unclear whether such enrichment simply reflects the high concentration of molecules involved in cell cycle control as essential in embryogenesis. It is unknown whether such chemistry could converge on a *bona fide* decision-making mechanism that influences embryonic fate, especially before it switches from a highly proliferative to a differentiating state at the onset of morphogenesis. We postulate that the conserved mtROS-based switch identified here could accomplish this by monitoring the redox status of embryos in a window preceding morphogenesis. As oxidants regulate key morphogenetic pathways involving PI3K/Akt, MAPKs, GPCRs and JNK signaling^101^, the development could be suspended until – and if at all – a sufficient redox capacity is achieved to promote morphogenesis.

Our work demonstrates in flies that limiting H_2_O_2_ levels can single-handedly regulate entry into such suspended animation by seizing embryonic contractility beyond a simple cell-cycle arrest. Such response is also embodied in zebrafish embryos, in which most cytoplasmic motion appears to be halted upon oxidant depletion (Fig. 4b and Supplementary Video 5), just as observed with anoxia-induced suspended animation in these embryos^5^. Readily hinted at by other recent studies^102,103^, these results may also provide a new rationale for why oocytes suppress ROS production: that H_2_O_2_ may otherwise act as an upstream chemical cue licensing morphogenesis. Further work is required to test whether this is mutually exclusive with the possibility of ROS as deteriorating agents in oocytes, as proposed before^100^.

What could sense declining H_2_O_2_ levels at the basis of an mtROS-based switch prior to morphogenesis? Unlike mechanisms that monitor oxidative stress, molecular mechanisms for sensing reductive stress are only beginning to be recognized^104,105^ and remain largely unexplored in animal development^106^. Although the suspended animation phenotype we observe here is essentially the same as that observed under anoxia conditions, mechanisms that monitor declining H_2_O_2_ levels are unlikely to converge on canonical oxygen sensors. Indeed, flies homozygous null for *sima* (*Drosophila* HIF-1α) are viable and fertile in normoxia^107^, and even the anoxia-response itself is independent of HIF-1α as demonstrated in worm embryos^6^. These suggest that a yet-to-be-identified mechanism in development must respond to declining H_2_O_2_ levels upon a wholesale depletion of either oxygen or oxidants.

We still do not know the conditions that activate this switch prior to the onset of morphogenesis and deactivate it thereafter. However, our study establishes the missing link between oxygen and morphogenetic licensing, invigorating future work to fully characterize this relationship. For instance, since O_2_^-^ molecules are detectable during blastoderm formation (Fig. 2a,b) at levels above basal production in normally functioning mitochondria, it is plausible that the ETC might be partially uncoupled prior to morphogenesis. A recent study reporting ETC uncoupling to gate the Spemann organizer in frogs attests to this possibility in a developmental setting^108^. In this regard, it is possible that cytoplasmic conditions favoring ETC uncoupling may help define when the mtROS switch can be active prior to morphogenesis.

By revealing a direct link between the early developmental utility of oxygen and an mtROS-based switch that controls morphogenetic commitment, our findings establish a mechanistic basis for uncovering the metabolic licensing of early embryogenesis. Further genetic and chemical studies dissecting the targets of such an mtROS-based response, and how declining H_2_O_2_ levels can be monitored in embryos, will pave the way for the molecular framework underlying this essential metabolic switch that controls the onset of animal morphogenesis.

## Acknowledgements

We thank Profs. Bruce Alberts and Shelagh Campbell, as well as members of the Aydogan Laboratory, for their support, insightful discussions, and critical read of this manuscript. To the benefit of humankind, yet to the curse of the authors, the field of redox metabolism is vast, and so is developmental biology: we apologize for any unintended omissions of bibliography, or those that we could not have meaningfully discussed due to space limitations. The research was funded by NIH DP2GM154328 (M.G.A.), NIH P30DK098722 UCSF NORC P&F Award (M.G.A.), Sandler Foundation Investigator Award (7029760; M.G.A.), and New Frontier Research Award by UCSF Program for Breakthrough Biomedical Research (7031159; M.G.A.). L-E.J. was supported by NIH R01GM144435. M.O. and O.D. were supported by a Swiss National Science Foundation Starting Grant (TMSGI3 218007).

## Author contributions

This study was conceptualised by K.M.T. and M.G.A. Investigation was done by U.K., F.D.R., S.J.P., L-E.J., O.D., K.M. and M.G.A. Data were analysed by U.K., F.D.R., S.J.P., M.O., L-E.J., O.D. and M.G.A. Methodology was developed by U.K., F.D.R., S.J.P., L-E.J., O.D., K.M. and M.G.A. Project was administrated by M.G.A. Resources were shared/made by U.K., F.D.R., S.J.P., M.O., K.M.T., L-E.J., O.D., K.M. and M.G.A. Software work was carried out by U.K., F.D.R., S.J.P., O.D. and M.G.A. Overall supervision was done by M.G.A. Validation experiments/analyses were carried out by U.K., F.D.R., S.J.P. and M.G.A. The main version of this manuscript was drafted by U.K., F.D.R., S.J.P. and M.G.A. with significant input from all authors. Finally, all authors reviewed and edited the manuscript.

## Data and material availability

All microscopy data and experimental materials are available from the corresponding author upon request.

## Methods

### *D. melanogaster* husbandry, embryo collection, and microinjections

Flies were kept at a 25°C incubator in culture medium (7.5% molasses, 1.01% agar, 1.4% agar, 5.6% cornmeal, 0.75% tegosept, 0.23% propionic acid, 0.04% phosphoric acid) in vials or bottles, as described previously^109^. All crosses were performed at room temperature and maintained at 25°C, unless mentioned otherwise. Following fly stocks were used in this study: OregonR wild type; His2Av-GFP (FBal0104781); His2Av-mRFP (FBst00233650); MRLC-GFP (FBal0221190); (sqh)-Moe-ABD-GFP (gift from D. Kiehart); UAS-Tom20-(SNAP)-GFP-(HA) (FBst0084254); UAS-Tom20-mCherry (gift from P.H. O’Farrell); V32-Gal4 (FBtp0009293); His2Av-mCer (FBst0091659).

For embryo collections, adult female flies were maintained in cages with juice plates (40% cranberry-raspberry juice containing 2% sucrose and 1.8% agar) supplemented with a droplet of yeast paste. To promote egg-laying and discard of over-night embryos, the cages were treated with a shedder plate for an hour prior to the embryo collection. Fresh juice plates were swapped with the shedder plate and incubated for 20 minutes at 25°C. The embryos were aged for an extra 45-50 minutes to aim for injections at nuclear cycle 9-11. Depending on the developmental cycle of interest, the incubation times varied. After the incubation period, embryos were dechorionated on a clear double-sided tape by manually removing the chorion using a needle and aligning them through a strip of glue on 35-mm MatTek glass-bottom petri dishes with a 14mm microwell, as described previously^110^. After desiccation at 25°C for 6 min, the embryos were covered with Grade H10S *Voltalef* oil (Arkema), followed by injections of drugs, dyes, purified enzymes or metabolites.

Microinjections were performed using Sutter Instrument borosilicate glass capillary tubes (of 1.2mm and 0.9 mm outer and inner diameters respectively). The capillaries were pulled on a P-87 micropipette puller at heat (670), pull (60), velocity (80), and time (190). Post-injection incubation times were 10 mins, except for purified enzymes injected and incubated 1h prior to imaging at desired cycles. Double injection experiments were performed by a first injection followed by a 10min incubation at 25°C, then a second injection aiming at the exact injection site (marked by the previous injection’s liquid displacement) followed by another 10min incubation at 25°C.

### *D. rerio* husbandry, embryo collections, and microinjections

Wild-type NHGRI-1 fish^111^ were bred and maintained using standard procedures^112^. Embryos were obtained by natural spawning and staged as previously described^113^. The researcher (Li-En Jao) was approved by the Institutional Animal Care and Use Committee, Office of Animal Welfare Assurance, University of California, Davis.

Microinjection of zebrafish embryos was performed as previously described^114^. Briefly, thin wall glass capillaries (World Precision Instruments, #TW100F-4) were pulled on a P-87 micropipette puller. Injections were performed with an air injection apparatus (Pneumatic MPPI-2 Pressure Injector). Injected volume was calibrated with a microruler. Approximately 2–3nl of an injection mix containing the drug or DMSO along with 48 μM 5-610CP-Hoechst^115^ was injected into the yolk of 1-cell stage embryos. Injected embryos were raised at 28.5°C until 512–1k cell stage before the major wave of zygotic genome activation^1^. The embryos were then manually dechorionated and mounted in 0.8% low-melt agarose with the dorsal side down in 35-mm glass-bottom dishes (MatTek Corp., P35G-1.5-10-C, or Cellvis, D35C4-20-1.5-N) for imaging.

### *S. arctica* culturing and maintenance

Cryopreserved in 2012 at −80°C, a frozen culture of the ichthyosporean *Sphaeroforma arctica* (originally described by Jøstensen et al.^116^) has been recently diluted and maintained in Marine Broth 2216 (MB) (BD Difco^TM^ #279110 or Sigma-Aldrich #76448, 37.4 g/L) at 17°C.

*S. arctica* cultures were grown and synchronized as described previously^91,94^. Briefly, saturated cultures in MB were diluted into fresh medium at a low density (1:200 dilution of the saturated culture) and cultivated in rectangular canted neck cell culture flasks fitted with vented caps (Falcon, #353108) at 17°C in the absence of light, leading to the development of a synchronized culture. Saturated cultures of *S. arctica* were obtained after ∼3 weeks of growth in MB or ∼5 days of growth in MB diluted to 1/16 with artificial seawater (Instant Ocean, 36 g/L).

### Administration of fluorescent probes, small molecules, purified enzymes and metabolites

#### Information on fluorescent probes

Following fluorescent probes with respective concentrations and solvents were microinjected into the indicated fly embryos: **Tetramethylrhodamine** (ThermoFisher, #T669) at 10µM in 100% DMSO into V32-Gal4; His2Av-Cer / UAS-Tom20-GFP embryos. **CellROX^TM^ Deep Red** (ThermoFisher, #C10422) at 250 µM in milliQ water into either wild-type or V32-Gal4; His2Av-Cer / UAS-Tom20-GFP embryos. **BioTracker Far-Red ONOO^-^ live cell dye** (Sigma-Aldrich, #SCT052) at 5mM in 100% DMSO into embryos expressing **a)** His2Av-GFP only, **b)** V32-Gal4; His2Av-Cer / UAS-Tom20-GFP embryos, or **c)** His2Av-GFP embryos pre-injected with Carboxy-PTIO potassium salt. **MitoPY1** (Tocris, #4428) at 1mM in 100% DMSO into His2Av-mRFP embryos. 100% DMSO used in these experiments was at 14M stock concentration as acquired from Fisher Scientific (#D136-1).

#### Information on small molecules

Following small molecules with respective concentrations and solvents were microinjected into the indicated fly embryos: **MitoTEMPO** (Sigma-Aldrich, #SML0737) at 5-10mM in 100% DMSO into wild-type embryos pre-injected with CellROX^TM^ Deep Red, or into embryos expressing **a)** His2Av-GFP, **b)** MRLC-GFP; His2Av-mRFP, or **c)** His2Av-mRFP; Moe-ABD-GFP. **S1QEL1.1** (Cayman Chemicals, #20982) at 1mM in 10% DMSO into either His2Av-GFP embryos, or wild-type embryos pre-injected with CellROX^TM^ Deep Red. **S3QEL2** (Cayman Chemicals, #18556) at 1.5 mM in 75% DMSO into His2Av-GFP embryos, or wild-type embryos pre-injected with CellROX^TM^ Deep Red. 100% DMSO was at 14M stock concentration as acquired from Sigma-Aldrich (#D8418). **Carboxy-PTIO potassium salt** (Sigma-Aldrich, #221) at 31.7mM in milliQ water into His2Av-GFP embryos. 100% DMSO used in these experiments was at 14M stock concentration as acquired from Fisher Scientific (#D136-1).

MitoTEMPO at 190-380mM in 100% DMSO was microinjected into zebrafish embryos bathed in 5mM MitoTEMPO (1-2% DMSO) throughout the experiments. *S. arctica* was cultured in 400µl media volumes containing 0.175mM effective concentration of MitoTEMPO.

#### Information on purified enzymes

Catalase from bovine liver (Sigma-Aldrich, #C1345) at 2,000-5,000 units/mg was dissolved in milliQ water to 0.5mg/ml and microinjected into fly embryos expressing His2Av-GFP.

#### Information on metabolites

Following metabolites with respective concentrations (at max.) and solvents were microinjected into the indicated fly embryos: **30% H_2_O_2_** (Fisher Scientific, #H325-500) at 9.8M in water into His2Av-GFP embryos. **ATP** (Fisher Scientific, #R0141) at 100mM in milliQ water into His2Av- GFP embryos. Starting with these max concentrations, serial dilutions and injections were performed for both metabolites to determine their effective physiological concentrations (see Fig. 3d and Extended Data Figs. 10 and 11). These solutions were then injected into His2Av-GFP embryos pre-injected with MitoTEMPO as part of the oxidant rescue experiments.

### Light microscopy

*D. melanogaster* embryos were live imaged alternatingly on three different spinning-disk confocal systems: PerkinElmer Ultraview using an Olympus IX70 microscope using with a planApo 60x 1.40 NA oil immersion objective; VT-QLC100 (VisiTech International) using a Leica DM-IRB microscope with a HCX PL APO 63x 1.40 NA oil immersion objective; CSU10 Yokogawa using a Nikon Eclipse Ti-E microscope with a perfect focus system and a 60x 1.4NA oil immersion objective. On the former system, image acquisition was controlled via the Volocity 6.3 software (PerkinElmer), whereas the latter two via the µmanager software. Depending on the experiments, a stack of 10-20µm was acquired with a step size ranging 0.25-0.5µm at 10s, 30s or 1min intervals. To conduct intensity-based measurements, the laser power was pre-measured using a PM100D digital optical power meter (ThorLabs) prior to every experiment and was maintained at a nearly constant W within each experiment.

*D. rerio* embryos were live imaged using Dragonfly (Andor Technology, Oxford Instruments), a spinning-disk confocal system using a Leica DMi8 inverted microscope with a HC PL APO 40x 1.10 NA water immersion objective. Image acquisition was controlled by Fusion software (Andor Technology), and images were captured every min using an iXon Ultra 888 EMCCD or Zyla sCMOS camera (Andor Technology). Time-lapse imaging was performed inside a wrap-around environmental incubator (Okolab) set at 28.5°C.

*S. arctica* were live imaged using a fully motorized Nikon Ti2-E epifluorescence inverted microscope, equipped with a hardware autofocus PFS4 system, a Lumencor SOLA SMII illumination system, and a Hamamatsu ORCA-spark Digital CMOS camera. Imaging was facilitated using CFI Plan Fluor objectives, including 20x (0.50 NA), 40x air, and 60x oil (0.5-1.25 NA). To maintain a constant temperature of 17°C during live microscopy, a cooling/heating P Lab-Tek S1 insert (PeCon GmbH) connected to a Lauda Loop 100 circulating water bath was employed. Fixed and immunostained *S. arctica* were imaged using an upright Leica SP8 confocal microscope, equipped with an HC PL APO 40x 1.25 NA glycerol objective.

### Electron microscopy

Sample preparation and electron microscopy experiments were done as previously described^117,118^. Briefly, wild-type embryos were collected at different developmental intervals and dechorionated by a 2min wash in 50% bleach. After a thorough rinse with water, the samples were transferred to the holder of a Balzer HPM 010 High Pressure Freezer, frozen and transferred to liquid nitrogen for long-term storage.

Samples were freeze-substituted in 2% osmium tetroxide plus 0.1% uranyl acetate in acetone for three days at −90°C. Subsequently, they were gradually warmed to 20°C over 6h and rinsed with 100% acetone for 3x 20min each. Infiltration with Epon-Araldite resin was carried out in a graded series of resin/acetone over 6 hours, then overnight in pure resin. Polymerization of resin was accomplished by heating at 60°C for 48 hours. 80nm sections were cut on a Reichert Ultracut E microtome and placed on copper grids.

Grids were stained with uranyl acetate for 5min and lead citrate for 2min. Using a Phillips CM10 electron microscope operating at 80kV, most micrographs were taken at a magnification of 15,500x. To generate the scale bar on EM micrographs in this paper, we took the diameter of microtubules (∼25nm) that are positioned longitudinally as our reference size.

### Ultrastructural Expansion Microscopy (U-ExM)

U-ExM on *S. arctica* was conducted as described previously^94,119^. Initially, cells were fixed using a 4% formaldehyde (FA) solution in 250mM Sorbitol. After two washes with 1x PBS, fixed cells were resuspended in 20-30μl PBS. These cells were then allowed to adhere onto 12 mm poly-l- lysine-coated coverslips for 1h. Subsequently, they were anchored in an AA/FA solution (1% Acrylamide (AA)/0.7% Formaldehyde (FA)) overnight at 37°C. For gelation, a monomer solution consisting of 19% (wt/wt%) sodium acrylate (Chem Cruz, AKSci #7446-81-3), 10% (wt/wt%) Acrylamide (Sigma-Aldrich #A4058), and 0.1% (wt/wt%) N, N’- methylenbisacrylamide (Sigma-Aldrich #M1533) in 1xPBS was used. The gels were polymerized for 1h at 37°C in a humidified chamber. To facilitate denaturation, the gels were transferred to a denaturation buffer (50 mM Tris pH 9.0, 200 mM NaCl, 200 mM SDS, adjusted to pH 9.0) for 15min at room temperature, followed by incubation at 95°C for 1h. Following denaturation, expansion was achieved through multiple water exchanges, following the previously established procedure. Post-expansion, the diameter of the gel was measured to determine the expansion factor. In our U-ExM images, scale bars represent the actual size, which is adjusted for the gel expansion factor.

### Sample fixation and immunostaining

*S. arctica* were fixed using 4% formaldehyde and 250mM Sorbitol for 20 min, then washed twice with PBS. For staining the nuclei, the cultures were allowed to settle for 15 min at room temperature before fixation. Subsequently, Hoechst 33342 (ThermoFisher, #62249) was added at 20 μM. Prior to imaging, fixed and stained *S. arctica* were concentrated and then mounted between a slide and a coverslip. For the U-ExM experiments, immunostaining of Histone H3 (Cell signaling, #9715) was performed using a 1/200 dilution followed with a 1/1000 diluted donkey anti-rabbit Alexa Fluor 488 secondary antibody (ThermoFisher, #A-21206). Antibodies were prepared in 3% PBS with 0.1% Tween 20.

### Image analysis and quantifications

All images were analyzed using ImageJ (v1.52), and all figure panels were assembled with Adobe Illustrator CC 2020. Graphs were generated using either GraphPad (Prism 10) or ggplot2 in *R* (v4.0.5.). Statistics analyses were performed using internal functions of GraphPad (Prism 10), and all relevant information on statistics tests are reported in respective figures or legends. Decision to perform the exact statistics tests were made based on whether the data were normally distributed by a D’Agostino & Pearson normality test.

#### Fluorescence signal quantification and normalization against mitochondrial mass in fly embryos

Following protocol was employed for all the TMRE, JC9, CellROX and MitoPY1 signal quantifications. Images were pre-processed in ImageJ (v1.52) and were cropped to consistently sized stacks with the injected areas at their centers where the fluorescent signal was at its maximum. The mean intensity of each slice was measured using *Measure* function in ImageJ and averaged across the z-stack. To measure the background for each image, four equidistant slices across the stack were taken and manually traced within a region of interest (ROI) around His2Av- marked nuclei, or around the nuclear “shadows” that emerged due to backlighting of the cytoplasmic signal if there was no nuclear marker used. The mean background was subtracted from the mean signal within each embryo.

To normalize fluorescence intensities against the mitochondrial mass within each cell cycle, we calculated the mean fluorescence signal of mitochondria in embryos transgenically expressing Tom20 fused to a fluorescent protein (GFP or mCherry), where all the probe injection experiments were performed. The background for these images were also calculated as described above. The probe signals were then normalized based on the calculated mitochondrial mass within each cell cycle. As the MitoPY1 and CellROX double injections (with MitoTEMPO, S1QEL1.1 and S3QEL2) were not performed in embryos expressing a mitochondrial marker, we used the normalization standards calculated from the TMRE solo injection experiments.

For ONOO^-^ dye signal quantifications, the above protocol was largely followed, except that the background was calculated via the far-red channel in unperturbed His2-GFP embryos. As ONOO^-^ signal was not mitochondrial (Extended Data Fig. 7b), mitochondrial mass normalization was not applied.

#### Signal co-localization analyses

Whether signal foci from fluorescent probes co-localize with the mitochondrial signal was assessed manually. Aggregates of fluorescent foci were identified, and their position was compared to the position of mitochondria. If majority of the aggregation had an overlap with a mitochondrion, the aggregate was counted as localized to the mitochondria. If most of the aggregates did not overlap or had no overlap with any mitochondria, they were counted as not localizing to the mitochondria.

#### Quantification of mitochondrial dynamics and morphology in fly embryos

Percentage occurrence of mitochondrial fission and fusion dynamics were quantified in live embryos expressing V37-Gal4>UAS-Tom20-mCherry. Based on a common rule across many cell types that individual mitochondria go through alternating fission-fusion cycles^120^, an initial assessment revealed that mitochondria in early fly embryos also follow this pattern with an approximate period of 80-160s per fission-to-fusion cycle or vice versa (during the 12^th^ nuclear division cycle). Taking the entire duration of nuclear cycle 12 into account (720-900s), the behavior of individual mitochondria was manually examined for 5 or 6 time points (with 120s intervals) across the cell cycle. The behavior of randomly chosen mitochondria were assessed at each time point until detecting five mitochondria that either fused or underwent fission, as not all mitochondria displayed membrane dynamics within our ±120s intervals (see Extended Data Fig. 1b, where ∼50% of the mitochondria are dynamic). Thus, on average, a total of ∼10 mitochondria were assessed for each time point per embryo. A constitutive separation or unity of mitochondria affirmed the behavior of fission or fusion respectively (see Fig. 1c). Temporal references on different phases of the cell cycle were determined by the exclusion of cytoplasmic fluorescent signal from the nucleoplasm (marking interphase), to which the signal would flood in when the nuclear envelope breaks (mitotic entry), and vice versa, when it reforms (mitotic exit).

Upon examination of their dynamics, the same mitochondria were utilized for size measurements, using the *Line* function in ImageJ. As most mitochondria are relatively small and positioned straight in the blastoderm (Fig. 1c), a line was drawn from one end to the next for each mitochondrion to manually measure their sizes.

#### Guidelines on assessing mitochondrial dynamics and autophagy in electron micrographs

To assess EM snapshots of mitochondrial membrane dynamics and motility, as well as of autophagy, previous guidelines were directly followed^121–125^. Briefly, mitochondrial fusion was judged by the connection of two mitochondria “bulges” where cristae structures are positioned orthogonally with respect, or unparallel, to each other. Mitochondrial fission was judged by pinching of a mitochondrion where the two pinched bulges appeared to have a unidirectional, continuous cristae structure. Mitochondrial motility was inferred from bent or curved mitochondria. Existence of autophagosomes was judged by membranous structures engulfing large vesicles or other organelles. Autolysosomes were evident by the former description along with a filling of undigested lipids.

#### Phenotypic classification and quantitation of various karyotypes

In embryos transgenically expressing His2Av fused to a fluorescent protein (GFP, RFP or CFP), the karyotypes observed upon a variety of perturbations in this study were classified into several groups based on the following criteria: **Normal**, when nuclei were like their counterparts during interphase or mitosis in unperturbed wild-type embryos. **Telophase defects**, when nuclei were arrested at a stage when their wild-type counterparts would be in telophase. **Chromosome damages**, when condensed chromosomes appeared broken and/or unevenly positioned in mitotic nuclei. The occasional nuclear fallouts observed during catalase injection experiments were classified as chromosome damages per prior literature^126^. **Desiccated**, when interphase nuclei appeared shrunken like peanuts, with sizes considerably smaller than expected average. **Rabl-like**, when chromosomes were condensed and lined against the inner wall of the nuclear envelope with all centromeres and telomeres at opposite poles of the nuclei – as described previously^3^. **Stippled chromatin**, when chromatin appeared in discrete conglomerates/speckles, but differently from Rabl-like, without an organized dispersal along the nuclear envelope. **Nuclear clumping**, when two or more nuclei were conjoined by their envelopes as though their centrosomes failed to separate. **Chromatin dispersal along NE**, when condensed chromosomes were lined-up against the inner wall of the nuclear envelope (NE) just as the Rabl-like phenotype, yet without a clear organization of the ends of chromosomes unlike it. This phenotype appeared to be a mixture of stippled chromatin and nuclear clumping phenotypes, as it displayed the features of both simultaneously. **Swollen and oblong**, when interphase nuclei appeared unusually large and/or in a non-round fashion. **Mitotic catastrophe**^126^, when mitotic chromatin appeared unusually disorganized in embryos attempting to cycle. **Firework**, when a portion of chromosomes were spread outward in a manner resembling this shape.

Care was taken to assign any observed phenotype – without omissions – into above classifications upon a variety of perturbations in this study. In each perturbed embryo, one of the above karyotype classes was observed as the defining phenotype (usually representing >80% of the nuclei in the blastoderm). Embryos were omitted from analysis only when they appeared **a)** sick, **b)** with sparse and uneven cytoplasm due to a harsh injection, **c)** with nuclei displaying high levels of karyotypic pleiotropy per the above classifications, or **d)** with a karyotype that failed to occur at a frequency that represent a statistically significant class per the above guidelines.

#### Quantifications of arrested *D. rerio* embryos

Zebrafish embryos injected with DMSO and MitoTEMPO were categorized into three time frames based on their survival into hours post-fertilization (hpf): 2-3, 3-4, and 4-5hpf. The embryos arrested in a particular time frame were not re-counted in the next time frame. The percentage of the ratio of surviving embryos were quantified and plotted.

#### Quantifications of nuclear content and division times in *S. arctica* cultures

For nuclear content distribution, the number of nuclei in fixed and Hoechst-stained *S. arctica* were counted using *ObjectJ* plugin in ImageJ. To compute nuclear division times, log_2_ of the geometric mean of the nuclear content was calculated as: log_2_(geommean) = ∑*_i_ f_i_* ∗ log_2_(*X_i_*) where f_i_ is the fraction of coenocytes and x_i_ the nuclear content (number of nuclei per cell) of each i^th^ nuclear content bin. All experiments were performed for a minimum of three independent times.

## Extended Data Figures

**Extended Data Fig. 1:**
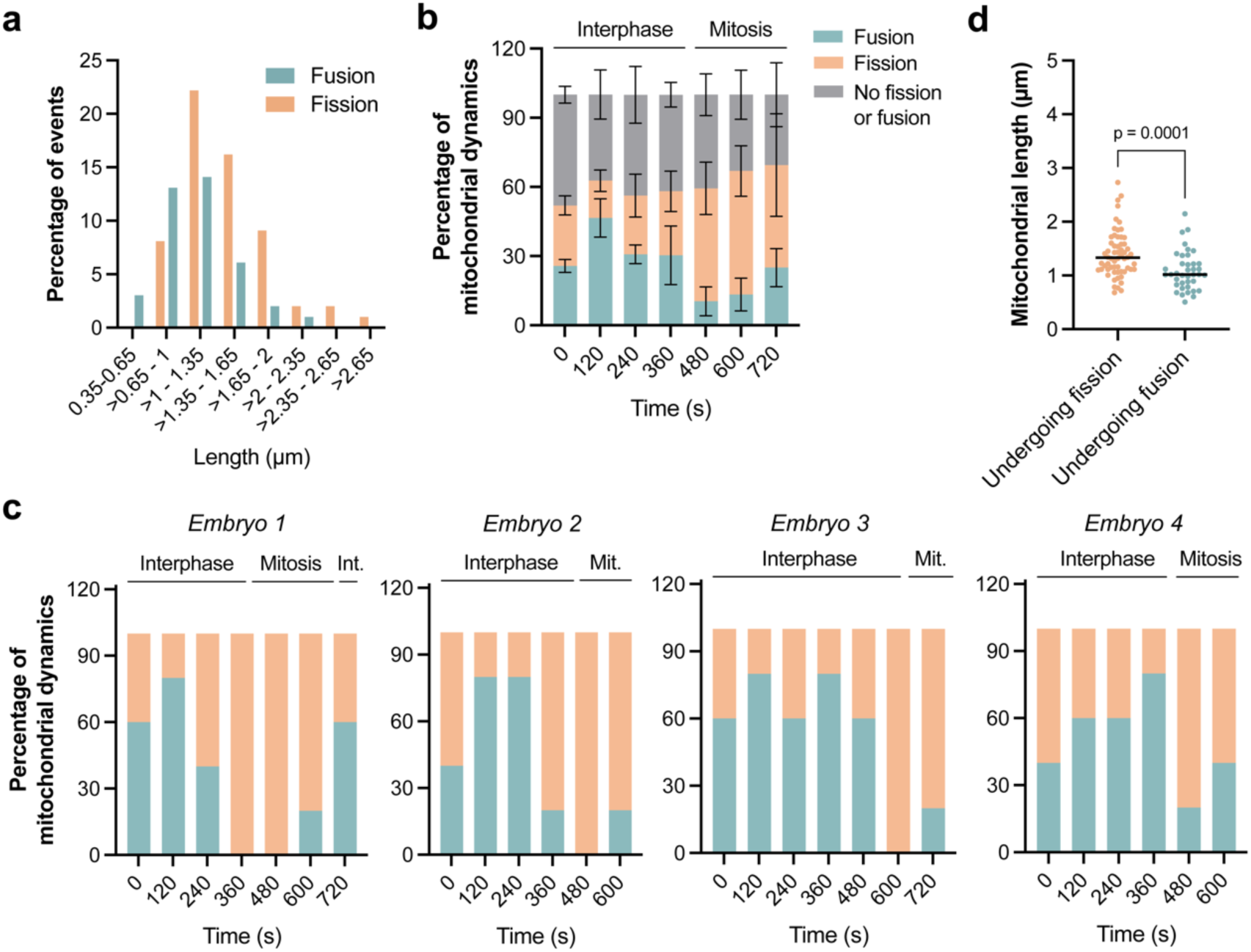
Characterization of mitochondrial morphology and dynamics during blastoderm formation in fly embryos. **a**, Bar graph distributing mitochondrial sizes associated with the percentage of fusion/fission events observed during our analyses in (**b** and **c**). **b**, Stacked bar graph quantifying the percentage of fusion or fission incidents observed for all mitochondria examined during cycle 12 (n>32 mitochondria per time point, except n=15 at t=720s). Data (mean ± s.e.m.) were plotted to represent an average of mitochondrial behaviour from multiple embryos. **c**, Same as in Fig. 1d except in individual embryos. **d**, Dot plots comparing the average length of mitochondria that fuse or undergo fission, using the data associated with (**b**). Data (mean ± s.d.) are compared using a Welch’s t-test.

**Extended Data Fig. 2:**
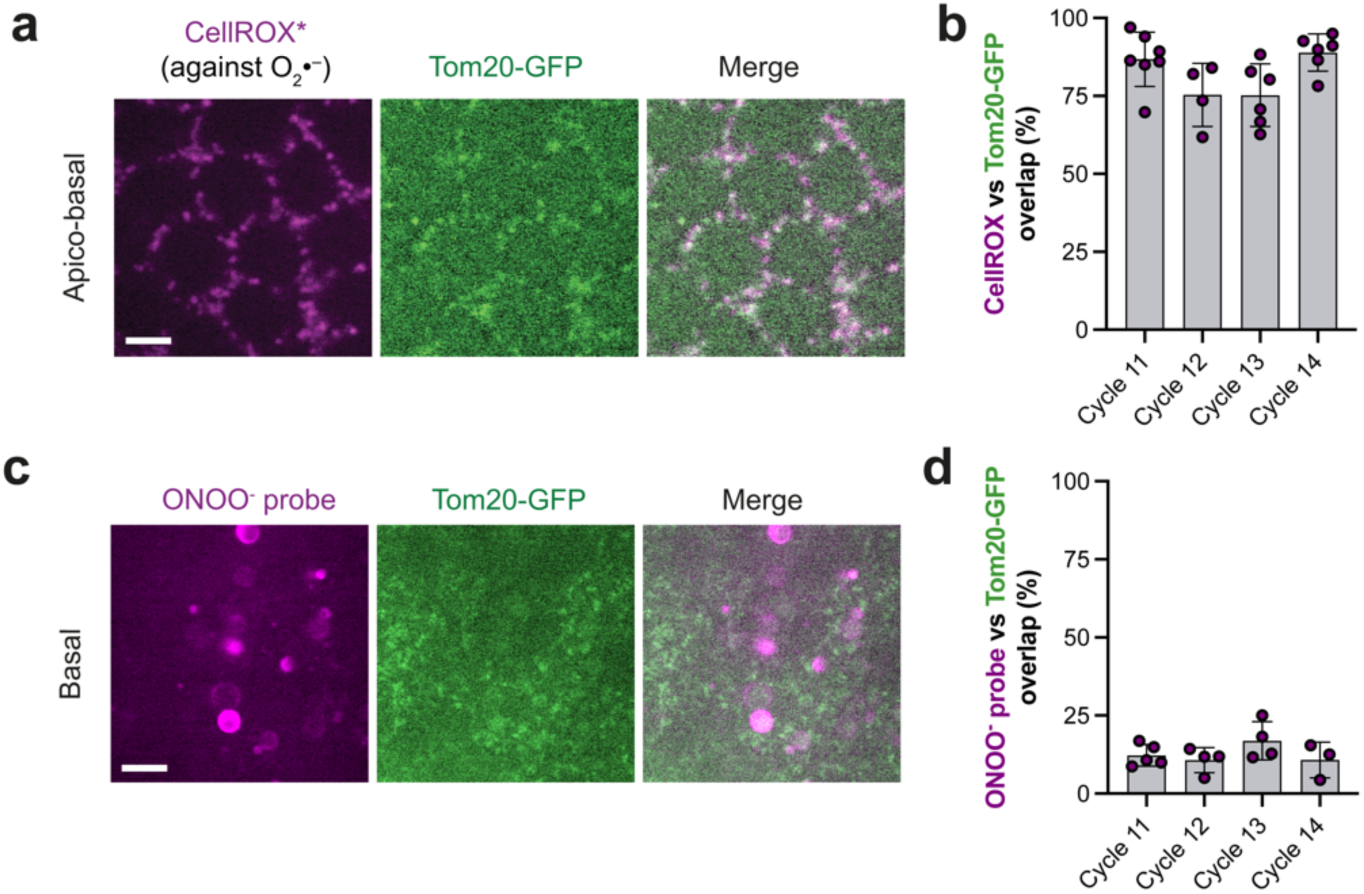
Signal overlap analysis for O_2_^-^ and ONOO^-^ probes against a mitochondrial marker. **a,** Micrographs depicting representative localization of O_2_^-^ in embryos expressing Gal4^V37^>UAS-Tom20-GFP injected with the CellROX probe. **b**, Bars quantifying the percentage overlap between the CellROX signal and Tom20-GFP mitochondrial marker as depicted in (a) (see *Methods*). Data (n≥4 embryos per nuclear cycle) are represented mean ± s.d., where each data point is the average of an embryo with a total of n≥1008 CellROX foci examined per nuclear cycle. **c**, Same as in a except for ONOO^-^ localization as detected by the ONOO^-^ probe. **d**, Same as in (b) except for quantifying the data in (c). Data (n≥4 embryos per nuclear cycle) are represented mean ± s.d., where each data point is the average of an embryo with a total of n≥248 ONOO^-^ probe foci examined per nuclear cycle. Scale bars, 5µm (**a** and **c**).

**Extended Data Fig. 3:**
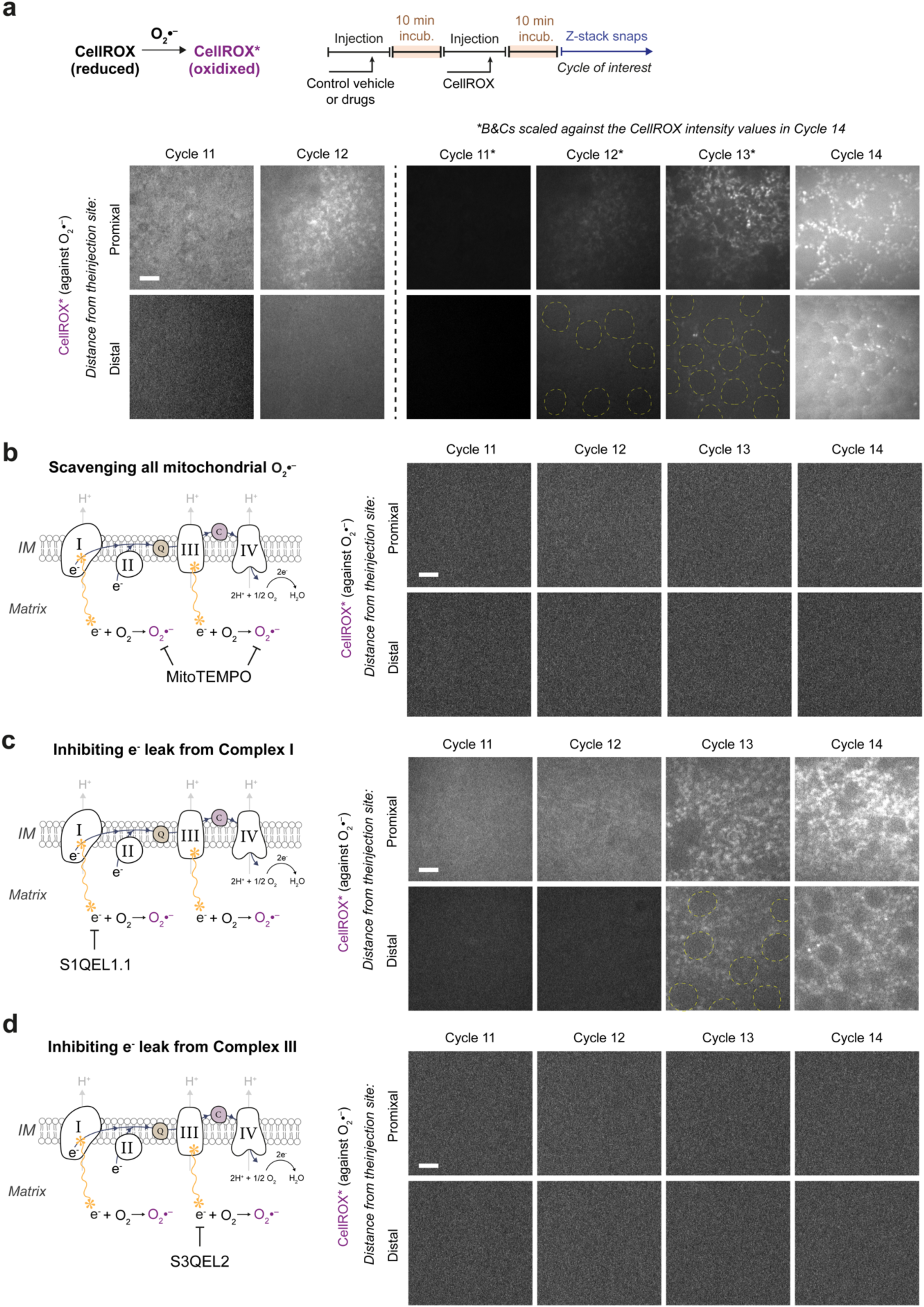
e^-^ leak from ETC Complex III can single-handedly account for the burst-like O_2_^-^ production during blastoderm formation. **a**-**d**, Micrographs showing CellROX probe signal in embryos administrated with (a) vehicle, (b) MitoTEMPO, (c) S1QEL1.1, and (d) S3QEL2. In (**a**), the brightness and contrast (B&C) of micrographs in cycle 11 and 12 were enhanced to demonstrate the basal level of O_2_^-^ production under an otherwise wild-type condition. B&C for the rest of the micrograph octets (**a**-**d**) were based on that of cycle 14 micrographs under control conditions in (a). *Proximal* and *Distal* indicates the proximity to the injection site. Given the fast turn-over of O_2_^-^, note the expected higher levels of CellROX in proximal regions. Enclosed *yellow* dotted lines in (a) and (c) signify nuclei. Sketch in (**a**) illustrates the experimental protocol (see *Methods*). Cartoons depict pharmaceutical strategies to (**b**) scavenge mitochondrial O_2_^-^ altogether or to (**c** and **d**) prevent e^-^ leak at prominent ETC sites that lead to O_2_^-^ production. Scale bars, 5µm (**a**-**d**).

**Extended Data Fig. 4:**
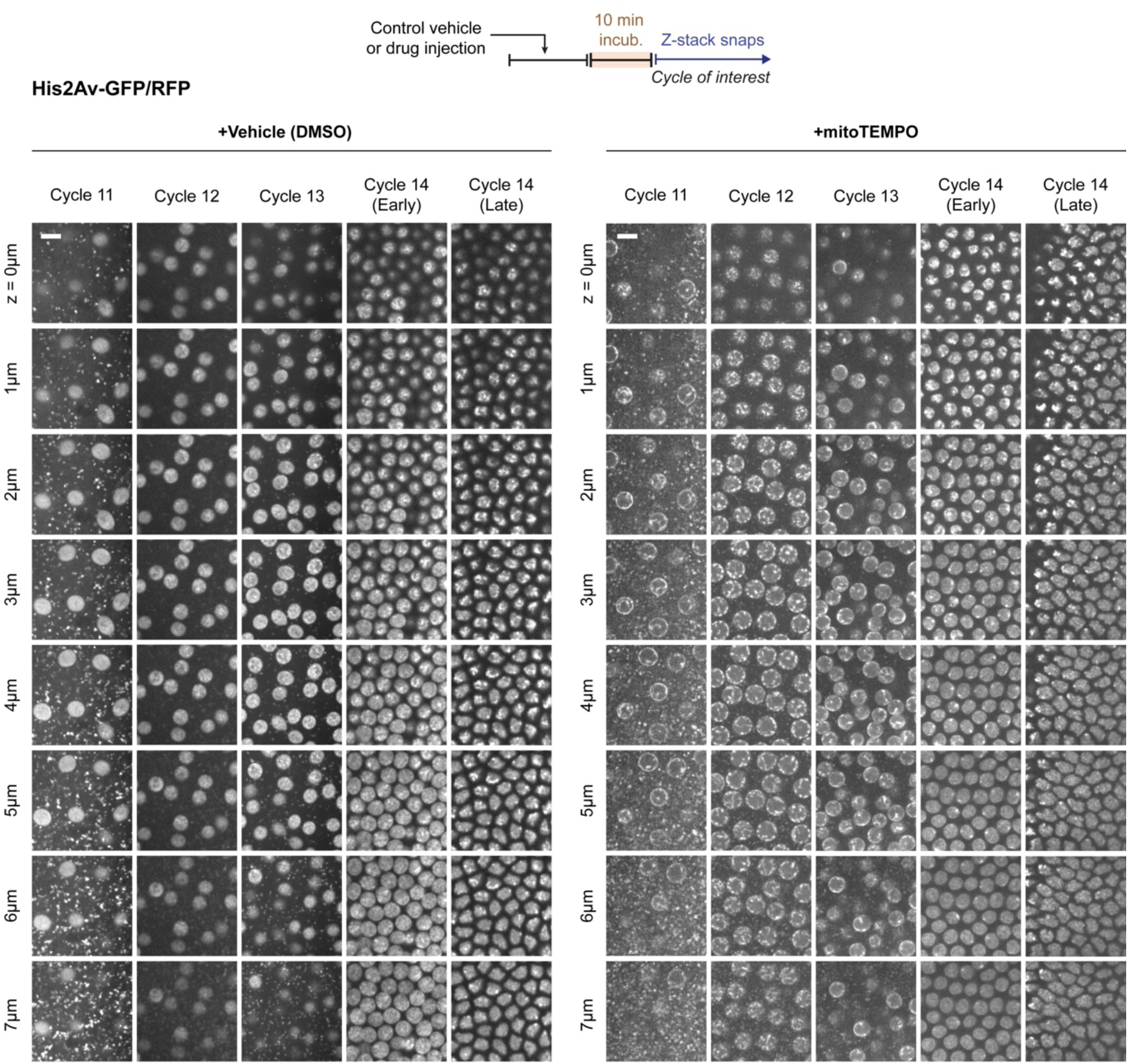
Scavenging mitochondrial O_2_^-^ altogether during fly blastoderm formation triggers a Rabl-like chromatin configuration à la anoxia in these embryos. Micrographs comparing the nuclear/chromatin phenotypes under control vs. MitoTEMPO injection conditions at cycles 11-14 in embryos expressing His2Av-GFP or -RFP. Note how euchromatin is normally largely diffuse and begins to condense into puncta-like regions through cycle 14 under control conditions. Meanwhile, MitoTEMPO treatment triggers a Rabl-like chromosome configuration à la anoxia in these embryos^3^: two ends of the chromosomes (telomeres/centromeres) are positioned across the apico-basal axis of nuclei, appreciated across z-depth slices. Sketch illustrates the experimental protocol (see *Methods*). Scale bars, 8µm.

**Extended Data Fig. 5:**
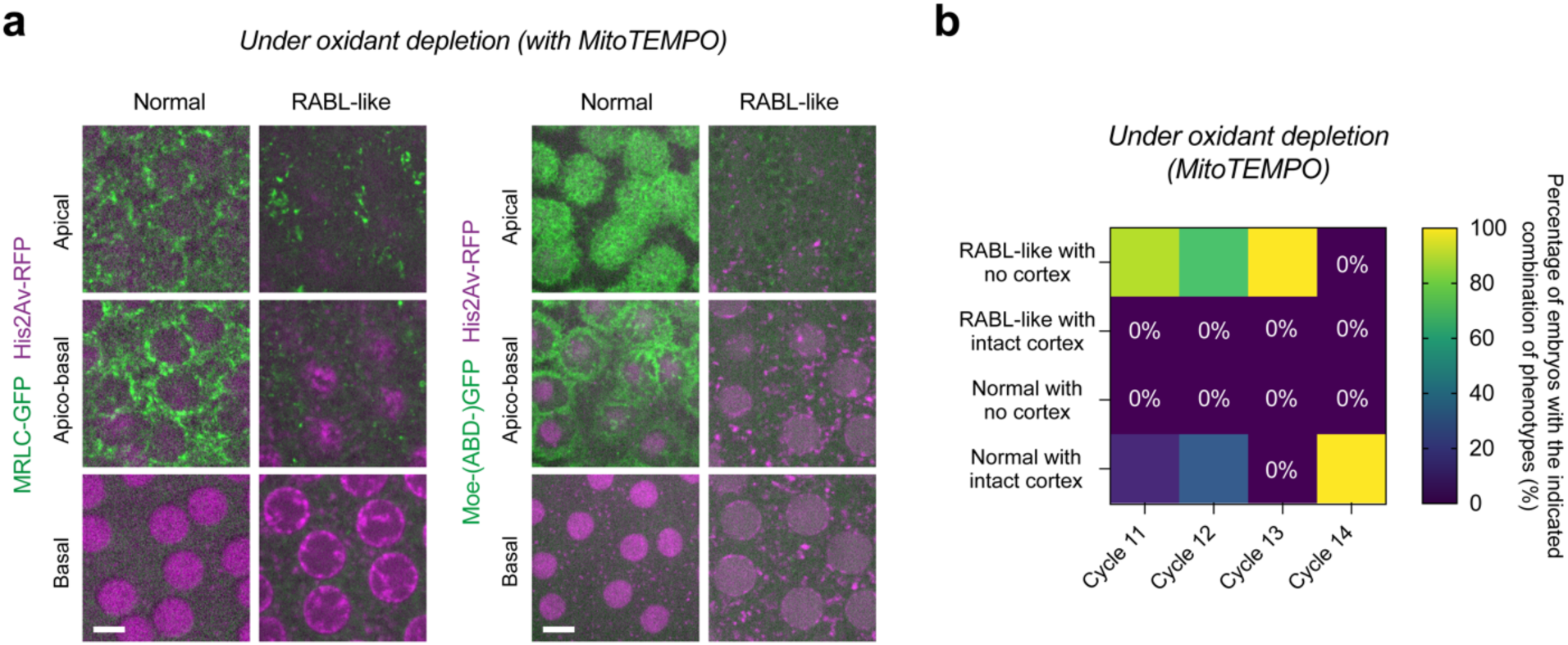
Oxidant-depletion triggered Rabl-like state is always associated with demolished cortical organization in the blastoderm. **a**, Micrographs from MitoTEMPO- injected embryos expressing His2Av-RFP along with either MRLC-GFP or Moe-(ABD-)GFP. Note the intact cortical organization when the nuclear morphology is normal, whereas it is disrupted entirely when the chromatin takes up the Rabl-like configuration in suspended animation. **b**, Heatmap quantifying the percentage of MitoTEMPO-injected embryos expressing His2Av-RFP and MRLC-GF from (a) with the indicated combination of phenotypes in blastoderms at cycles 11 (n=9), 12 (n=7), 13 (n=13) and 14 (n=11). Scale bars, 5µm.

**Extended Data Fig. 6:**
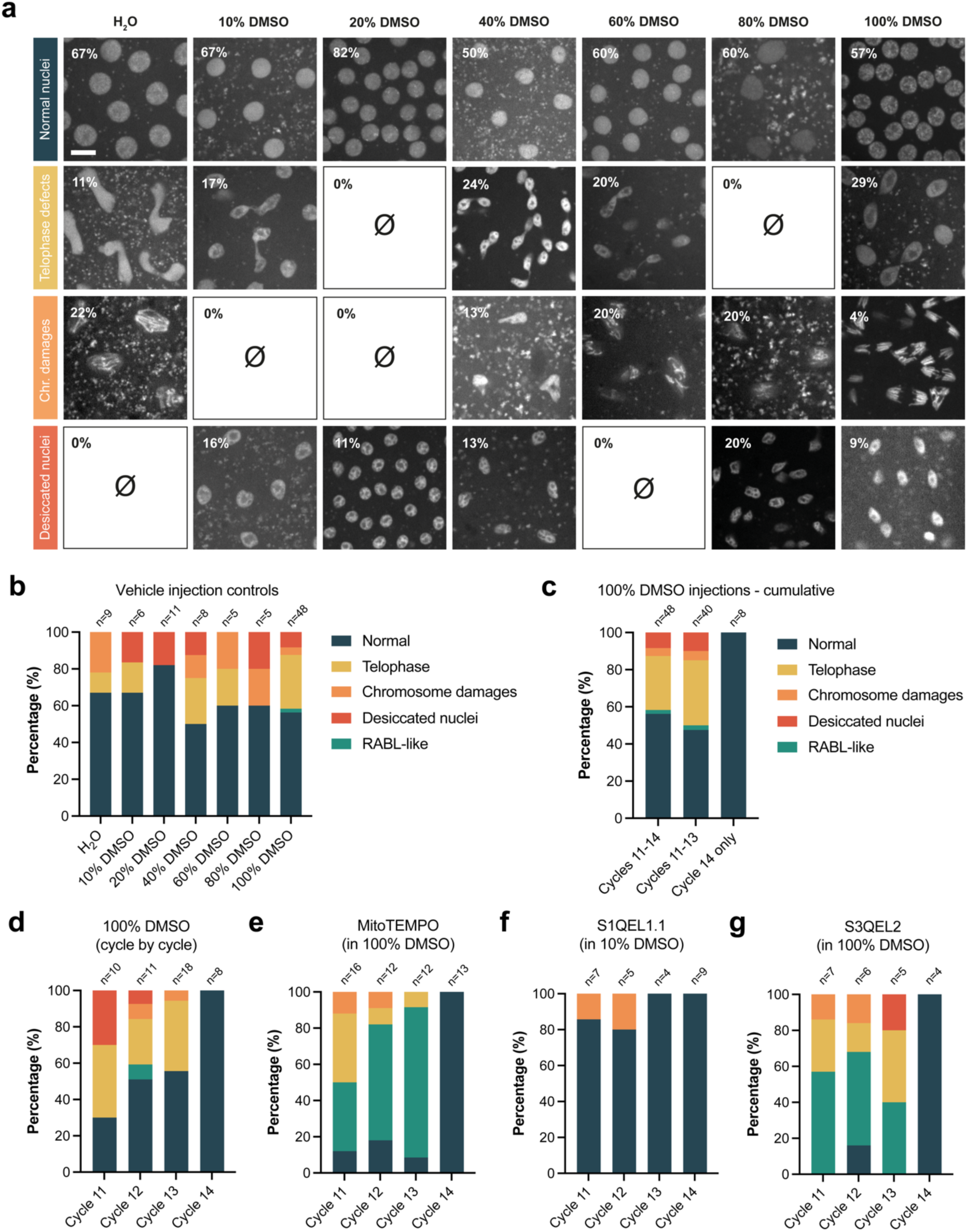
Nuclear and chromatin phenotypes observed in embryos treated with a dilution series of DMSO or with a series of pharmaceuticals that use DMSO as solvent. **a**, Micrographs demonstrating the gallery of colour-coded nuclear/chromatin phenotypes in His2Av-GFP embryos that were injected with the indicated dilution series of DMSO. Inset percentages signify the fraction of embryos with the associated phenotype under the indicated condition. **b**, Stacked bars quantifying the phenotypes from (a) with the associated sample (embryo) numbers for ease of comparison. **c**, Same as (b) except that it is only quantifying the 100% DMSO phenotypes collectively for cycles 11-13, cycles 11-14 or cycle 14 only. **d**-**g**, Stacked bars quantifying the cycle-by-cycle percentage distributions of the nuclear and chromatin phenotypes observed under (**d**) vehicle, (**e**) MitoTEMPO, (**f**) S1QEL1.1, and (**g**) S3QEL2 conditions. Scale bar, 8µm.

**Extended Data Fig. 7:**
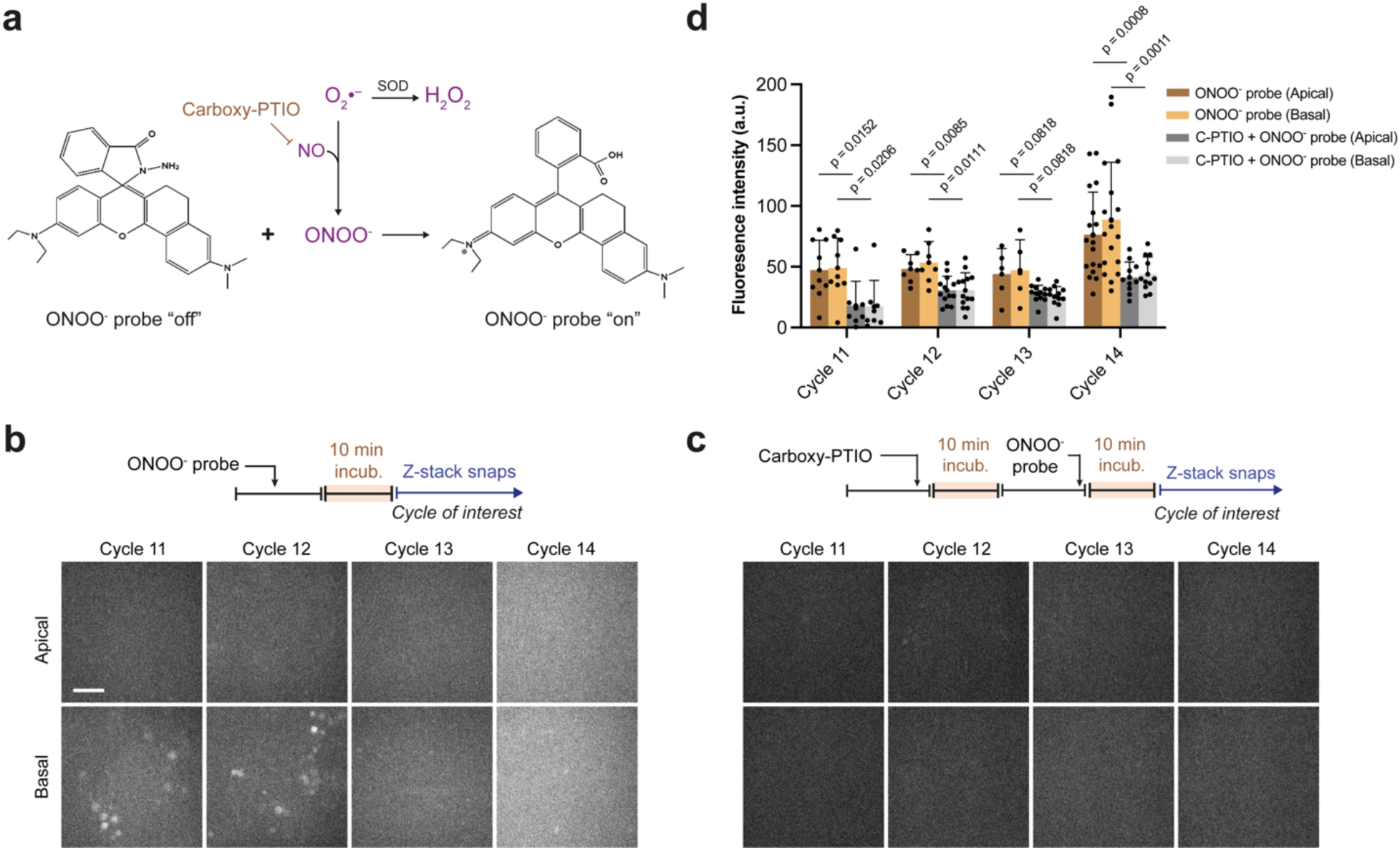
Scavenging NO can diminish levels of ONOO^-^ molecules, which first concentrate in yolk granules then largely diffuse in the cytosol during normal blastoderm formation. **a**, Reactions depicting the chemical landscape of reactive nitrogen species in relation to O_2_^-^ and its otherwise dismutation by superoxide dismutase (SOD). Note how the ONOO^-^ probe turns “on” to fluoresce by reacting with, hence consuming, ONOO^-^. **b**-**c**, Micrographs demonstrating the endogenous ONOO^-^ signal as detected by the injection of its probe into embryos either (**b**) solo or (**c**) after a pre-treatment with Carboxy-PTIO (C-PTIO), a potent NO scavenger. **d**, Bar graphs showing the significant reduction of ONOO^-^ levels in embryos treated with C-PTIO during various blastoderm cycles. Data (n≥5 embryos per nuclear cycle in each condition) are represented mean ± s.d. (where each data point represents an embryo), compared using a Welch’s t-test (for Gaussian distributed data) or a Mann-Whitney test. Sketches in (**b** and **c**) illustrate the experimental protocol (see *Methods*). Scale bar, 10µm.

**Extended Data Fig. 8:**
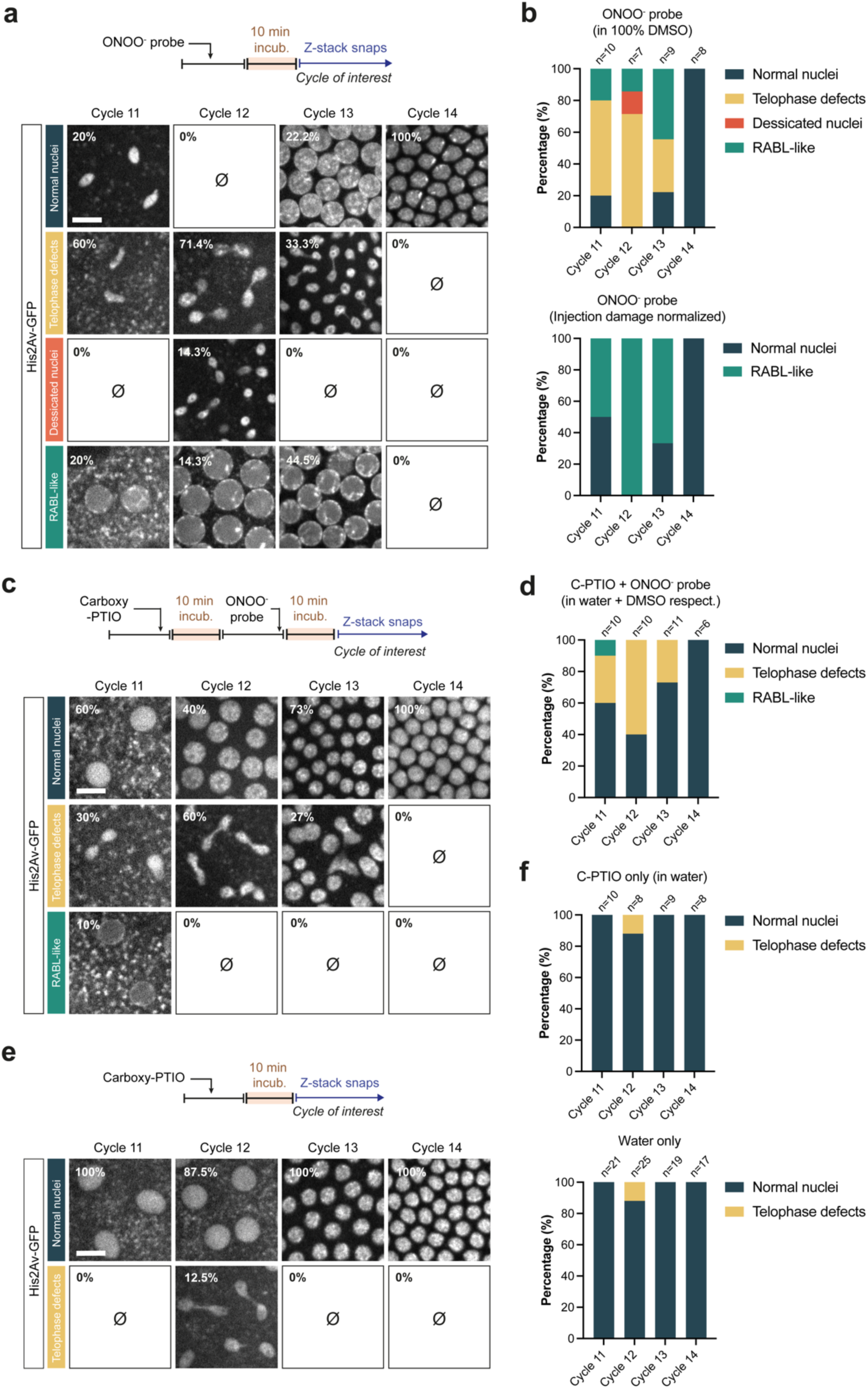
Rabl-like suspended animation in embryos injected with ONOO^-^ probe can be explained by diminishing O_2_^-^ levels. **a,** Micrographs demonstrating the gallery of colour-coded nuclear/chromatin phenotypes in His2Av-GFP embryos that were injected with the ONOO^-^ probe. **b**, Stacked bars (top) quantifying the phenotypes from (a) with their associated sample (embryo) numbers. To ease comparison between the normal vs Rabl-like conditions, stacked bars (below) quantify the same as above, except with the percentages normalized to exclude otherwise karyotype damages induced via the vehicle injection (100% DMSO) itself by default (see Extended Data Fig. 6d). **c**, Same as (a), except that the embryos were pre-treated with Carboxy-PTIO (dissolved in water) before the ONOO^-^ probe injection. **d**, Stacked bars quantifying the phenotypes from (c). **e**, Micrographs displaying the nuclear/chromatin phenotypes observed in His2Av-GFP embryos that were injected with Carboxy-PTIO. **f**, Stacked bar (top) quantifying the phenotypes from (e) with the associated sample (embryo) numbers. Stacked bars (below) quantifying phenotypes from the same experimental protocol that was performed for Carboxy-PTIO’s vehicle (water). Inset percentages (**a**, **c** and **e**) signify the fraction of embryos with the associated phenotype under the indicated condition. Sketches in (**a**, **c** and **e**) illustrate the experimental protocol (see *Methods*). Scale bars, 10µm.

**Extended Data Fig. 9:**
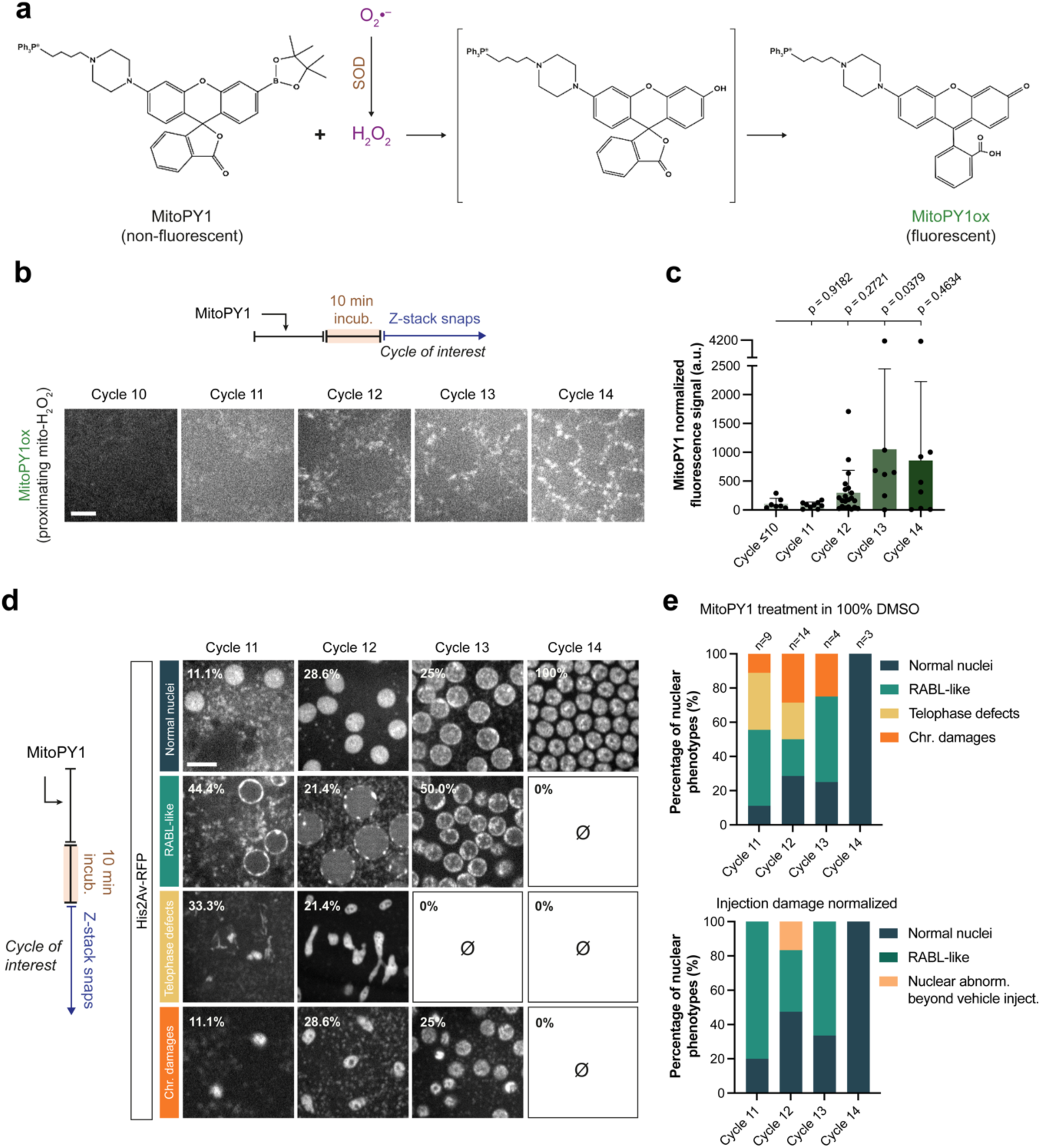
Evidence of mitochondrial H_2_O_2_ production and associated Rabl-like suspended animation upon its consumption during blastoderm formation. **a**, Reaction depicting how H_2_O_2_ is produced by superoxide dismutase (SOD) via the dismutation of O_2_^-^. Note how H_2_O_2_’s reaction with – hence consumption by – MitoPY1 triggers the removal of the boronate group from the latter and “turns on” its fluorescent properties by oxidation (signified by MitoPY1ox). As MitoPY1 contains a triphenyl phosphate moiety (Ph3P^⊕^), it enables the localized detection and imaging of H_2_O_2_ specifically in mitochondria. **b**, Micrographs demonstrating oxidized MitoPY1 signal (against H_2_O_2_) in wild-type embryos injected with this probe. **c**, Bar graph quantifying the mean MitoPY1 signal from the experiments (b) in cycle 10 (n=7), 11 (n=9), 12 (n=21), 13 (n=7), and 14 (n=8). Data are mean ± s.d. (where each data point represents an embryo), compared using a Mann-Whitney test. **d**, Micrographs displaying the nuclear/chromatin phenotypes observed in His2Av-RFP embryos that were injected with MitoPY1. Inset percentages signify the fraction of embryos with the associated phenotype under the indicated condition. **e**, Stacked bars (top) quantifying the phenotypes from (d) with the associated sample (embryo) numbers. To ease comparison between the normal vs Rabl-like conditions, stacked bars (below) quantify the same as above, except with the percentages normalized to exclude otherwise karyotype damages induced via the vehicle injection (100% DMSO) itself by default (see Extended Data Fig. 6d). Sketches (in **b** and **d**) illustrate the experimental protocols. Scale bar, 5um (**b**), 10µm (**d**).

**Extended Data Fig. 10:**
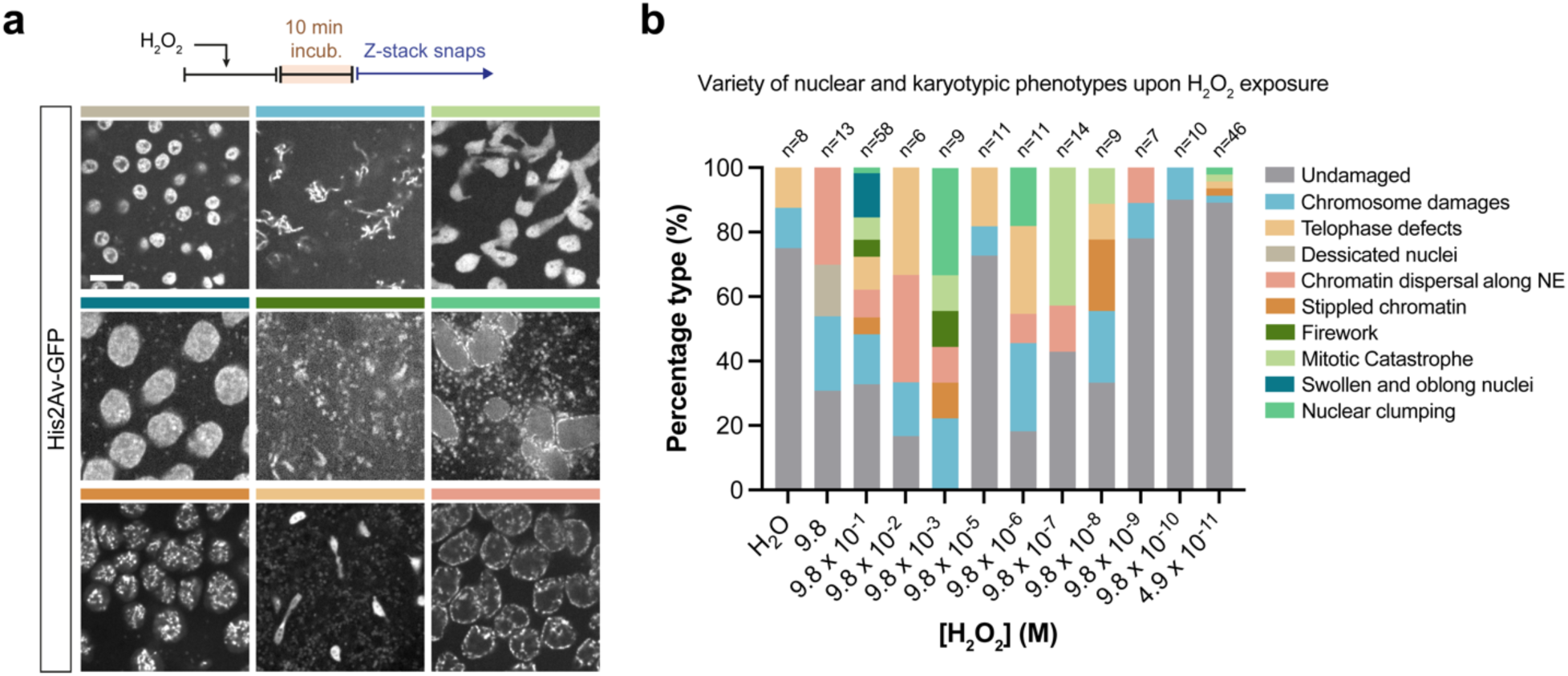
Gallery of karyotype damages observed in response to the injection of a dilution series of [H_2_O_2_] during blastoderm formation. **a**, Micrographs depicting the colour-coded nuclear karyotype damages associated with the injection of a dilution series of [H_2_O_2_] during blastoderm formation. See (b) for the legend of these phenotypes. Sketch illustrates the experimental protocol. **b**, Colour-coded stacked bars quantifying the fraction of embryos displaying the phenotypes from (a) as a function of varying [H_2_O_2_] administered. Associated sample (embryo) numbers are indicated above each stacked bar. Experiments with [H_2_O_2_] at 9.8×10^-1^ and 4.9×10^-11^ M were oversampled to ensure whether either of the extreme (deci- vs nano-molar) concentration ranges are indeed associated with as much (or as little) of damages as displayed by the consecutive concentrations. Scale bar, 5µm.

**Extended Data Fig. 11:**
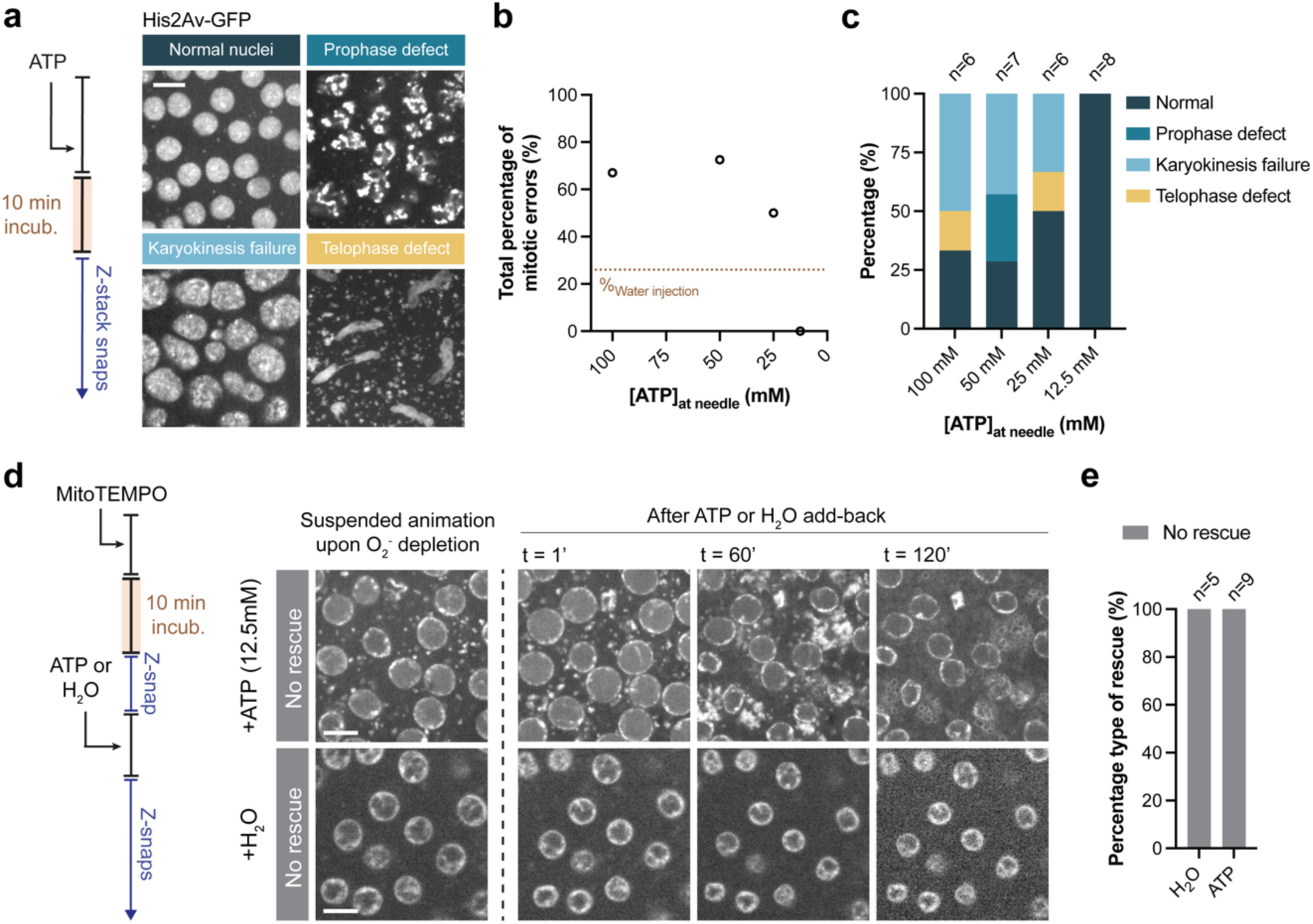
Neither ATP nor water can rescue oxidant depleted embryos from suspended animation. **a**, Micrographs demonstrating representative physiological nuclear (normal) or pathological mitotic karyotypes associated with the injection of a dilution series of [ATP] into embryos expressing His2Av-GFP. **b**, Scatter plot quantifying the percentage of embryos with the mitotic karyotype damages from (a) as a function of [ATP] with which His2Av- GFP embryos were treated. *Brown* dotted line represent the maximum level of specimen percentile with general karyotype damages even when injected with water (see Extended Data Fig. 6a). **c**, Stacked bar graphs quantifying the distribution of the mitotic error phenotypes from (a) observed as a function of ATP dilution series. Associated sample (embryo) numbers are indicated above each stacked bar. **d**, Micrographs depicting His2Av-GFP embryos that were initially treated with MitoTEMPO to induce suspended animation and subsequently were injected with either ATP (12.5mM) or water to attempt to rescue them from this developmental arrest. **e**, Bar graphs quantifying the rescue attempts with ATP and water from (d) in MitoTEMPO-arrested embryos. Associated sample (embryo) numbers are indicated above each stacked bar. Sketches in (**a** and **d**) illustrate the experimental protocols. Scale bars, 5µm (**a** and **d**).

**Extended Data Fig. 12:**
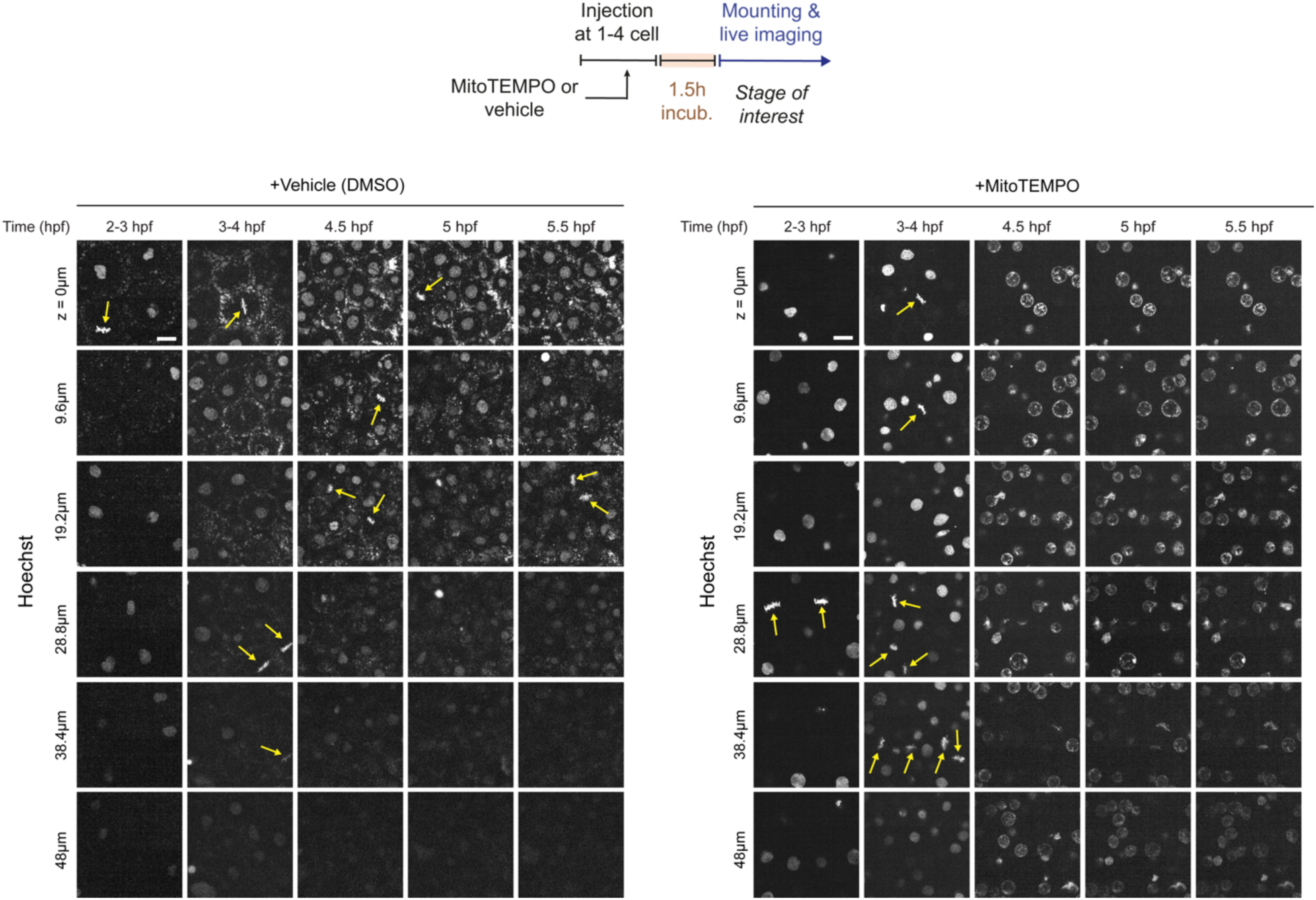
Scavenging mitochondrial O_2_^-^ in early *D. rerio* embryos triggers suspended animation à la anoxia in these embryos. Micrographs comparing the progression of cell division cycles under control vs. MitoTEMPO injection conditions in living *D. rerio* embryos pre-incubated with solution containing Hoechst (marking chromatin). Note the suspended animation with a distinct chromatin configuration in MitoTEMPO-injected embryos after 3-4 hours post fertilization (hpf), when any observable movement including cell divisions cease – phenotypes that directly mimic the effects of anoxia on these embryos^5^ (see Supplementary Video 5). *Yellow* arrows signify metaphase events indicative of cell division, or the lack thereof, e.g., post 3-4hpf under MitoTEMPO conditions. Sketch illustrates the experimental protocol. Scale bars, 20µm.

**Extended Data Fig. 13:**
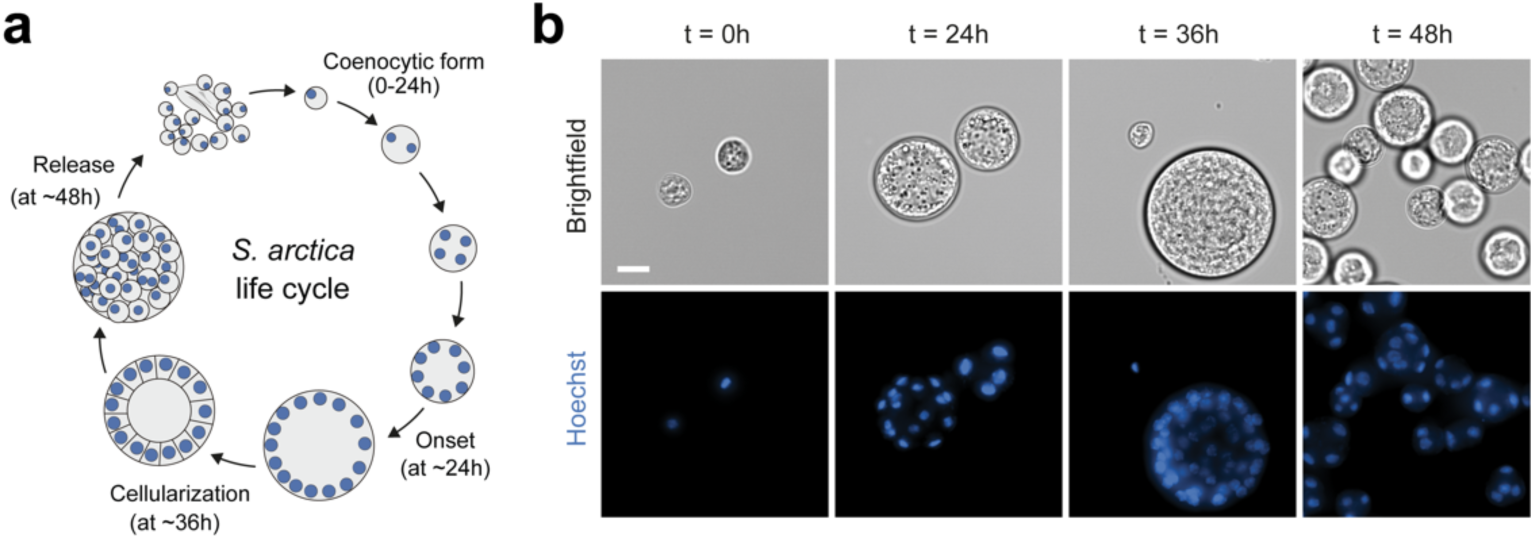
*Sphaeroforma arctica* (*S. arctica*) undergoes a morphogenetic event to cellularize and disperse into individual cells after a coenocytic developmental cycle. **a**, Cartoon depicting the life-cycle of close animal relative *S. arctica*, resembling the path to blastoderm formation in early animal embryogenesis^89^. **b**, Micrographs displaying representative points in the life-cycle of *S. arctica* fixed and stained with Hoechst (marking nuclei). Note the dispersal of cellularized *S. arctica* from 36 to 48h transition. Scale bar, 25µm.

## Supplementary Videos

Supplementary Video 1: Mitochondrial dynamics during blastoderm formation in flies expressing Tom20-mCherry.

Supplementary Video 2: mtROS depletion-triggered suspended animation in fly embryos (first half) with ‘no rescue’ phenotype upon H_2_O_2_ supplementation (4.9×10^-10^ M).

Supplementary Video 3: mtROS depletion-triggered suspended animation in fly embryos (first half) with ‘full rescue’ phenotype upon H_2_O_2_ supplementation (4.9×10^-10^ M).

Supplementary Video 4: mtROS depletion-triggered suspended animation in fly embryos (first half) with ‘partial rescue’ phenotype upon H_2_O_2_ supplementation (9.8×10^-10^ M).

Supplementary Video 5: mtROS depletion-triggered suspended animation in early zebrafish embryos.

Supplementary Video 6: mtROS depletion-triggered suspended animation in *Sphaeroforma arctica*.

